# Senescent tenocytes drive an age-associated loss of cell-extracellular matrix homeostasis, which can be partially restored using dihydroresveratrol

**DOI:** 10.64898/2026.07.26.740761

**Authors:** Jack Llewellyn, Nodoka Iwasaki, Anna Hoyle, Iolanda Vendrell, Georgina Berridge, Kathryn Collins, Hollie Delo, Roger K Smith, Jayesh Dudhia, Richard G A Faragher, Chavaunne T Thorpe

**Author notes:** Corresponding author: Correspondence to: Dr Chavaunne Thorpe.

## Abstract

Tendons are commonly injured, not only in athletes, but also during normal everyday activities, particularly in older age. Tendons are rich in extracellular matrix proteins, many of which are turned over extremely slowly during an individual’s lifetime. As a result, tight regulation of the extracellular matrix is essential for tendons to remain resilient to the mechanical load placed on them. However, tendons are prone to age-associated functional decline, characterised by chronic inflammation, accumulation of damaged collagen, and matrix remodelling. These degenerative changes often precede injury, wherein additional inflammation and fibrotic tissue deposition make treatment difficult and reinjury highly likely. Understanding and preventing the causes of tendon functional decline is therefore vital to improving quality of life in the ageing population.

Using an equine model, here we show that tendon fibroblasts, or *tenocytes*, isolated from aged tendons exhibit markers of senescence when cultured *in vitro*. Further, we describe how senescence in tenocytes drives inflammation, hypercontractility, and dysregulation of tendon matrix components. By replicatively senescing tenocytes isolated from young tendons, we observed that senescent tenocytes instigated proinflammatory signalling, had impaired capacity in wound healing assays, increased contractility, and secreted factors that induced senescence in healthy tenocytes. Further, tenocyte senescence dysregulated matrix turnover both at the gene and protein level. Finally, we demonstrate how senotherapeutic treatment of senescent tenocytes can reduce expression of senescence markers and restore proliferative capacity.

Our results uncover systemic links between tendon ageing and cellular senescence, and identify mechanisms and therapeutic strategies for age-associated tendon degeneration.

## Introduction

Tendons connect muscles to bone and are essential for movement. Specific tendons that are adapted for high-speed locomotion are highly elastic and have an energy storing function to improve locomotion efficiency (Biewener, 1998). Energy storing tendons are particularly prone to injury, both in athletes and the general population, with ageing a major risk factor for tendinopathy, which costs the European economy €115 billion/year, not only in treatment costs, but also due to the indirect costs of both work absence and reduced health due to an inability to exercise (Abbah et al., 2014).

Tendon injuries are also common in horses, with the equine energy storing forelimb superficial digital flexor tendon (SDFT) providing an excellent model to study tendon injuries, as the risks and characteristics of injury are remarkably similar to the human Achilles tendon (Lui et al., 2011; Patterson-Kane and Rich, 2014). Tendon injuries are recalcitrant to treatment; a poor understanding of the underlying molecular causes of tendinopathy across species means that there is a lack of effective treatments, poor functional healing, and a high rate of reinjury (Innes and Clegg, 2010; Patterson-Kane and Rich, 2014). In the horse, the reinjury rate is estimated to be 40-60 %, and up to 90 % of flat racehorses do not return to racing following injury (Smith, 2024). In humans, Achilles tendon degeneration generally precedes rupture (Hess, 2010). One study of patients over 65 presenting with Achilles tendon rupture found only 36 % returned to previous activity and 25 % experienced re-rupture within 3 years (Nestorson et al., 2000).

Chronic inflammation and formation of fibrotic tendon scar tissue, rich in glycosaminoglycans and type III collagen, underline this high likelihood of reinjury (Smith, 2024). Similarly, low-grade inflammation develops during ageing in a number of tissues (Franceschi et al., 2018) and has been linked to chronic tendinopathy (Rees et al., 2014). Further, tendon ECM composition is altered with ageing, in both horses and humans, concomitant with increasing incidence of tendon injury with age (Ely et al., 2009; Kotsifaki et al., 2026). There is therefore a pressing need to develop more effective treatments and preventative therapies for age-related tendon degeneration.

Cellular senescence is a hallmark of ageing (López-Otín et al., 2023) which has become increasingly implicated in musculoskeletal pathology (Wan et al., 2021) and ageing-related diseases (Childs et al., 2017; Raffaele and Vinciguerra, 2022), and its role in tendon degeneration is now beginning to emerge. Senescent cells lose their ability to grow and divide, and the senescence-associated secretory phenotype (SASP) can affect neighbouring cells, induce inflammation, and affect wound healing (Hernandez-Segura et al., 2018). In the shoulder, expression of senescence markers is significantly increased in the tendon tissue of patients with chronic tendinopathy (Bühler et al., 2025, 2022), and also as a consequence of corticosteroid treatment (Poulsen et al., 2014), while in human and rat Achilles tendon there has been significant progress in establishing the effect of ageing on tendon stem/progenitor cells (TSPCs), with studies demonstrating that TSPCs become senescent with advancing age (Kohler et al., 2013; Rui et al., 2019; Zhou et al., 2010). However, the effect of ageing on other tendon cell populations, including tenocytes, the predominant cell type within tendon, is not well characterised. While our previous work shows that tenocytes within the ageing SDFT upregulate genes associated with senescence and the SASP (Zamboulis et al., 2024), the consequences of the accumulation of senescent tenocytes in aged tendon, and methods to mitigate tenocyte senescence remain to be established.

Pharmacological interventions targeting senescent cells, referred to as senotherapeutics, are an emerging branch of regenerative medicines that have yielded beneficial effects in preclinical models of renal dysfunction, metabolic disorders, cardiovascular diseases, musculoskeletal impairment, respiratory disorders, central nervous system diseases, and skin conditions (Raffaele and Vinciguerra, 2022). While nomenclature is inconsistent, the increasingly common convention is to classify senotherapeutics as either senolytics, which selectively eliminate or target senescent cells for clearance; senoreverters, which revert the phenotypes of senescent cells to that of healthy cells; and senomorphics, which suppress the proinflammatory SASP of senescent cells (Park and Shin, 2022; Popescu et al., 2023; Wang et al., 2025). Herein we report the efficacy of candidate senotherapeutics of each of these classifications in the treatment of senescent tenocytes.

The most well-characterised senolytic treatment, Quercetin and dasatinib (Q+D) in cocktail, targets Bcl-xL and Src tyrosine kinase pathways typically upregulated in senescent cells (Anerillas et al., 2022; Mas-Bargues et al., 2021; Novais et al., 2026; Rivera-Torres and San José, 2019), and has been shown to selectively remove senescent human rotator cuff tendon cells (Hawthorne et al., 2024). Quercetin alone is a commercially available dietary supplement and improves injury repair in aged murine Achilles (Wang et al., 2026). Similarly, resveratrol is a commercially available dose-dependent SIRT1 activator which in turn inhibits the senescence marker p53 (Brockmueller et al., 2023), and has been shown to have protective and reparative effects in murine Achilles tendon healing (Da Ré Silva et al., 2021; Zeytin et al., 2020) and in equine tenocyte cultures (Heidari et al., 2024). Resveratrol and its primary metabolite dihydroresveratrol (dRSV) have also been shown to act as senoreverters, augmenting telomerase activity and allowing senescent cells to re-enter the cell cycle (Latorre et al., 2017; Wang et al., 2011), with downstream effects including improved resilience to oxidative stress, increased cell proliferation, and decreased expression of inflammatory signalling pathways (Li et al., 2025). However, crucially, dietarily administered resveratrol is rapidly metabolised *in vivo* with almost no unmetabolised resveratrol detected in human plasma, versus micromolar concentrations of the metabolite (dRSV) (Faragher et al., 2011; Walle et al., 2004), and thus we opted for the metabolite as our candidate senoreverter. Finally, SB203580 (international nonproprietary name adezmapimod (Adz), is a p38 MAPK inhibitor which has shown promise in lowering expression of SASP components at clinical trial (Alimbetov et al., 2016; Hongo et al., 2017). Further, it has been shown to be beneficial to murine rotator cuff (Wilde et al., 2016) and Achilles tendon repair (Schwartz et al., 2015).

In this study, we characterise the consequences of senescence in equine tenocytes, and develop methods to rescue this process. We find multiple markers of senescence in tenocytes isolated from aged tendons, and establish a replicative senescence model to link phenotypes of tenocyte senescence to symptoms of tendon degeneration. Using this model, we report senescent tenocytes as proinflammatory and contractile, with impaired wound healing capacity. Additionally, these cells dysregulate tendon matrix synthesis and turnover, and secrete factors which induce senescence in healthy tenocytes. We then utilise our model to characterise the effects of an array of senotherapeutics. Each compound trialled improved the transcriptomic similarity between senescent and healthy cells, with dRSV in particular restoring proliferation and alleviating markers of senescence, outlining the potential of senotherapeutics as a novel regenerative medicine against tendon degeneration.

## Results

### Markers of senescence present in tenocytes from aged tendon

Senescence was confirmed in tenocytes shortly after isolation from aged SDFT by positive staining of the lysosomal enzyme β-galactosidase (Figure 1A), a canonical marker of senescent cells owing to increased lysosomal content of the enzyme (Kurz et al., 2000). Further, tenocytes isolated from aged tendons exhibited slower growth *in vitro* relative to tenocytes isolated from young tendons, with an average doubling time of 54 hours versus 35 hours (p = 0.0134, Figure 1B). Finally, tenocyte cultures isolated from aged tendons also had more than double the proportion of cells expressing the cyclin-dependent kinase inhibitor p21 (3.5 % versus 1.5 %, p = 0.0429, Figure 1C), indicative of a greater proportion of the cell population having undergone the permanent cell cycle arrest associated with cellular senescence. These results, taken together with previous data, indicate an accumulation of senescent tenocytes in aged SDFTs.

**Figure 1.**
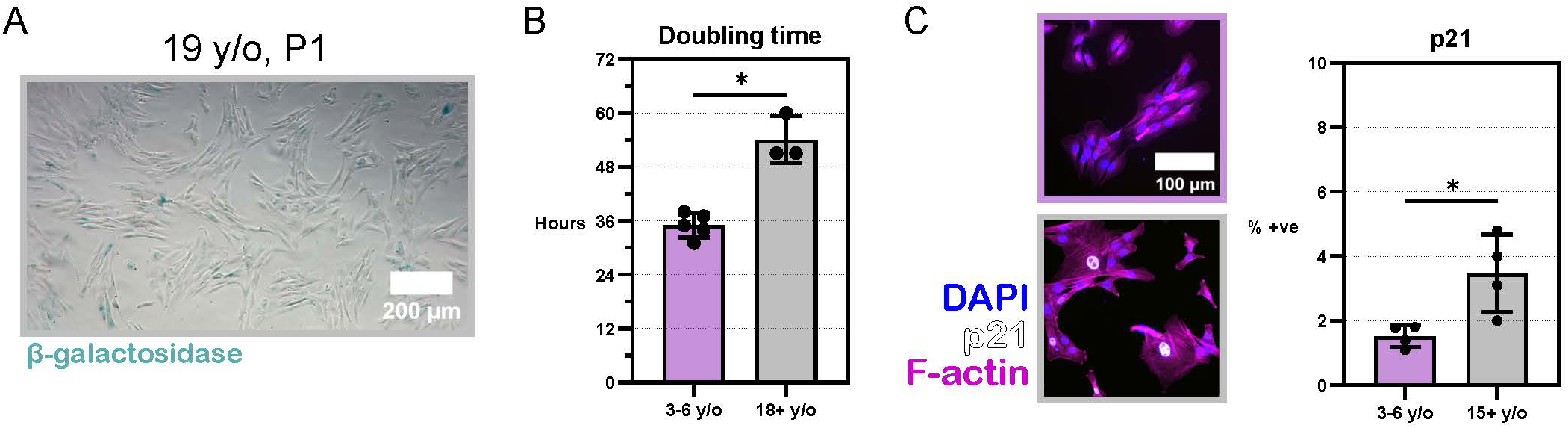
Markers of senescence present in tenocytes isolated from aged equine superficial digital flexor tendons (SDFT). **(A)** Tenocytes isolated from aged SDFT and stained for senescence-associated β-galactosidase at passage 1. **(B)** *In vitro* doubling times of tenocytes isolated from either young (purple) or aged (grey) SDFT. N = 5 (young), 4 (aged). **(C)** Immunofluorescence of p21 protein in tenocytes isolated from either young (purple) or aged (grey) SDFT. N = 4. Data are represented as mean ± SD, and p values show the results of a Welch’s unpaired two-tailed t test (∗ < 0.05).

### Repeated passaging induces senescence in equine tenocytes

To decipher the functional consequences of increasing numbers of senescent cells in ageing tendon, we sought to develop an *in vitro* model of tenocyte senescence. Replicative senescence is of *in vivo* relevance to cells reaching their Hayflick limit in geriatric organisms and, as opposed to other methods such as those utilising radiation exposure or oxidative stress, yields a more translationally relevant mixed population of healthy and senescent cells (Herr et al., 2024; Veronesi et al., 2023). Tenocytes isolated from the healthy SDFTs of young, skeletally mature horses aged 2-6 years were divided into two populations. For each biological replicate, one population of cells was cryogenically stored as a low-passage (Lo-P) control, while another population of cells was repeatedly passaged in culture over the course of several months, and designated as high-passage (Hi-P) at the point of clear changes in growth and morphology, before validation using canonical markers. Figure 2A demonstrates a consistent doubling time of tenocytes through passaging before a gradual slowing of growth above passage 11. Our previous data indicates the increase in doubling time is due to an increasing portion of the population entering growth arrest, rather than an elongation of the cell cycle (Kalashnik et al., 2000). Passages were consistently defined in relation to seeding density, culture media, and confluency; and a 72-day period of recording population doublings (Supplemental Figure 1B) demonstrated the growth of each replicate plateauing, suggesting tenocyte populations isolated from young, skeletally mature equine SDFT generally are capable of more than 40 doublings.

**Figure 2.**
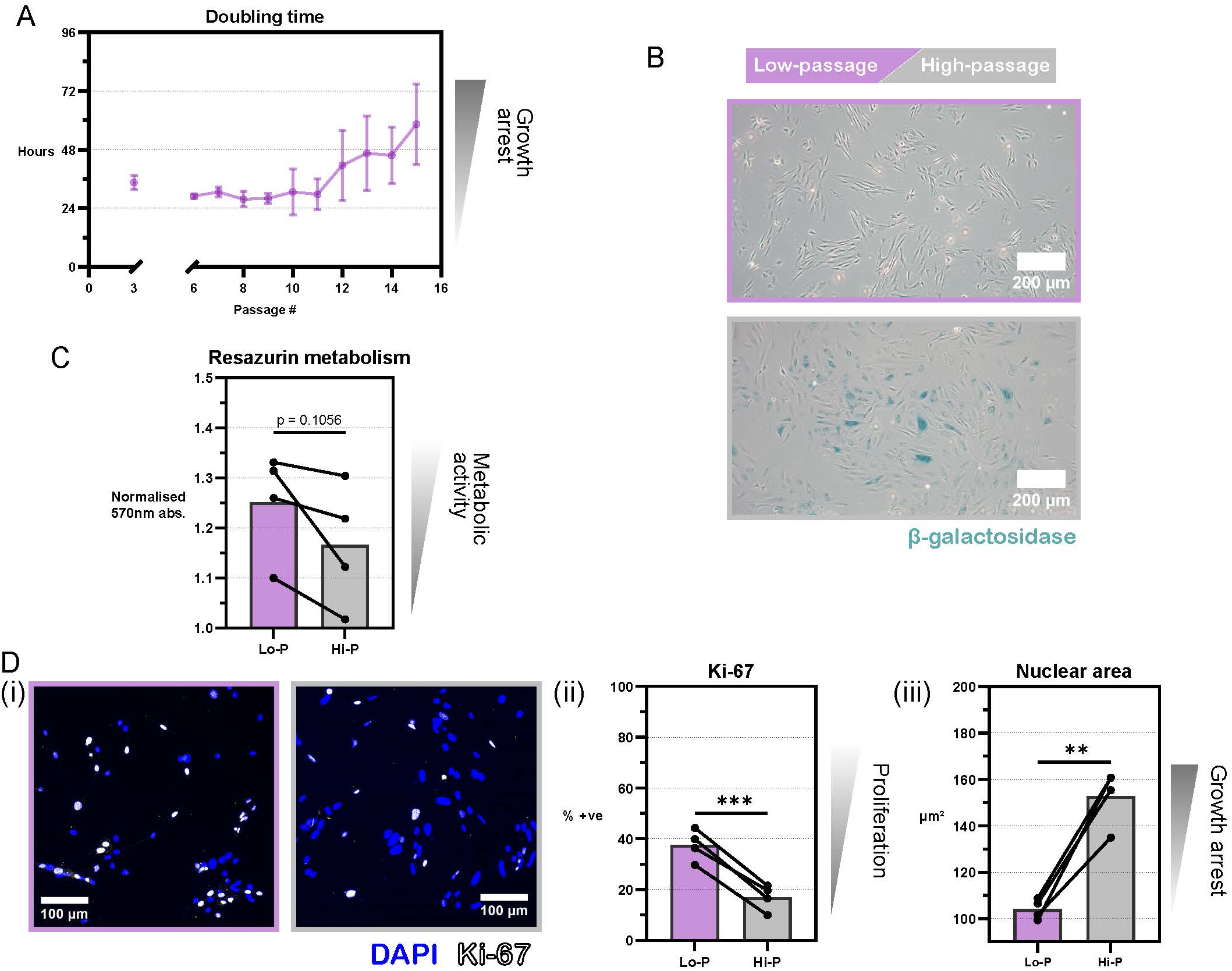
Repeated passaging induces senescence in equine tenocytes. **(A)** *In vitro* doubling times of tenocytes isolated from young, healthy superficial digital flexor tendons, senesced through repeated passaging. Data are represented as mean ± SD, N ≥ 3 for each passage #. **(B)** Naïve (low-passage (Lo-P), ≤ P5, purple) and replicatively stressed (high-passage (Hi-P), ≥ P10, grey) tenocytes, stained for senescence-associated β-galactosidase. N = 4, see supplementary replicate images. **(C)** Blank-normalised 570nm absorbance of Lo-P and Hi-P cells treated with 200 µM resazurin to quantify cellular metabolism. N = 4. **(D)** (i,ii): immunofluorescence of S-phase marker Ki-67 protein in tenocytes isolated from either Lo-P or Hi-P tenocytes. (iii): quantification of nuclei size in immunofluorescence images of Lo-P and Hi-P cells. N = 4. Unless otherwise specified, bars represent means of biological replicates, and p values show the results of a paired two-tailed t test (∗∗ < 0.01, ∗∗∗ < 0.001).

This method generated biological replicate-paired populations of Lo-P and Hi-P tenocytes, which allow us to infer the direct consequences of senescence in tenocytes with fewer confounding factors than comparing cells from young and aged tendons. Senescence in Hi-P populations was confirmed using several canonical markers. Positive β-galactosidase expression was observed in Hi-P populations (Figure 2B, Supplemental Figure 1A), concomitant with reduced metabolic activity in all replicates, as observed through a minor decrease in metabolism of resazurin supplemented into the culture medium (Figure 2C, p = 0.1056). Hi-P populations also contained far fewer cells expressing the proliferation marker protein Ki-67 (Figure 2D, p = 0.0009), which is highly upregulated in cycling cells (Miller et al., 2018). We also observed changes in nuclear morphology, shown to be a cross-species marker of senescence (Heckenbach et al., 2022), with Hi-P tenocytes having an enlarged average nuclear area as viewed from above using 2D microscopy (153 µm² versus 104 µm² in Lo- P cells, p = 0.0035), accompanied by a slight decrease in the expression of the nuclear structural protein Lamin B1 (Supplemental Figure 1C). Senescence induced via administration of 200 nM dexamethasone demonstrated that increased lysosomal activity (Supplemental Figure 1D) and cell cycle arrest markers (Supplemental Figure 1E) are phenotypes of tenocyte senescence which are conserved across senescence-triggering stimuli.

### RNA sequencing identifies activation of inflammatory pathways and dysregulation of tendon matrix proteins by senescent cell populations

Changes in tenocyte gene expression driven by cellular senescence were characterised by RNA sequencing subject to false discovery rate (FDR) corrections, comparing Lo-P and Hi-P populations. Importantly, significant downregulation of just one tendon marker gene was observed as a result of extended passaging *in vitro* (thrombospondin 4, Supplemental Figure 2A). Of relevance, thrombospondin 4 is known to decrease with age in mice (Boisvert et al., 2018; Gan and Südhof, 2019). Hi-P expression of type I collagen, type III collagen, scleraxis, and tenascin C did not significantly decrease compared to Lo-P cell levels, while tenomodulin was not expressed in *in vitro* tenocytes of any passage, as has been reported previously (Jo et al., 2019).

We identified 11 % of 17898 transcripts with significantly altered expression after false discovery rate (FDR) correction in Hi-P populations, and through Gene Set Enrichment Analysis (GSEA) we detected significant activation/suppression in 9 of the 50 Hallmark gene sets after FDR correction (Liberzon et al., 2015) (Figure 3A-B). Further validation of an accumulation of senescent cells in Hi- P populations was also seen, with the CDK inhibitors p16 and p21 both significantly upregulated in Hi-P cells (log_2_ fold-changes 4.00, 1.50 and FDR-adjusted p values 1.75×10^-18^, 2.95×10^-4^ respectively, Figure 3B). The activated gene sets in Hi-P populations consisted of the proinflammatory TNFα signalling via NFκB and Inflammatory response pathways, typically associated with the SASP. As seen in other fibroblast cell types (Llewellyn et al., 2023), metabolic and proliferation pathways featured heavily among the suppressed gene sets in Hi-P populations, consisting of Bile acid metabolism, Cholesterol homeostasis, Fatty acid metabolism, Xenobiotic metabolism, E2F targets, G2/M checkpoint, and mTORC1 signalling.

**Figure 3.**
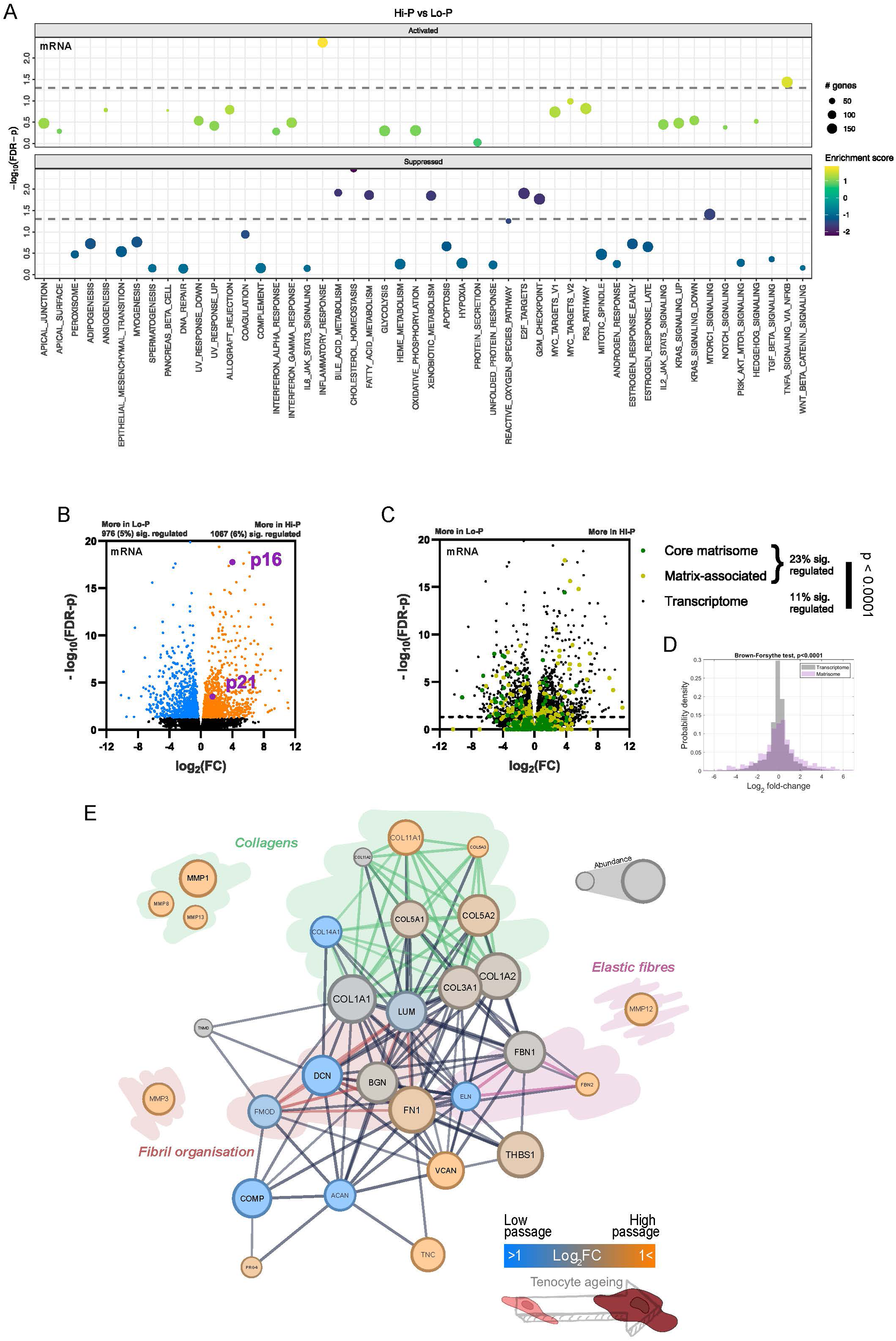
RNA sequencing characterises dysregulation of the tendon extracellular matrix by senescent tenocytes. **(A)** Gene Set Enrichment Analysis (GSEA) of enrichment in the Hallmark gene sets between low- (Lo-P) and high-passage (Hi-P) tenocytes. Dashed line indicates a false discovery rate-corrected p value < 0.05 (Subramanian et al., 2005). N = 4. **(B)** Volcano plot of transcriptional changes between Lo-P and Hi-P tenocytes. Coloured points satisfy a false detection rate-corrected p value < 0.05. Blue indicates significant downregulation in Hi-P, orange indicates significant upregulation in Hi-P, p values from RNA sequencing are calculated as described in Methods. **(C)** The same volcano plot shown in (B), with coloured points indicating transcripts corresponding to the matrisome (Shao et al., 2023). **(D)** Browne-Forsythe test comparing the distribution of fold changes of the transcriptome versus the matrisome. **(E)** Interaction network with nodes consisting of transcripts detected within our sequencing data whose proteins have known function in tendon. Nodes are sized according to their signal strength in low-passage cells in our sequencing data, coloured according to their fold-change in high passage cells, and edges are weighted by StringDB interaction score (version 12.0), showing only edges which satisfy the highest confidence filter.

Of particular note, several of the highest fold-changes and most significant changes in gene expression corresponded to the extracellular matrix (ECM). Figure 3C illustrates that while 23 % of the matrisome (Shao et al., 2023) showed significantly differential gene expression in Hi-P populations, versus 11 % of the transcriptome as a whole, expression of ECM genes was not consistently lower or higher in Hi-P populations., This indicates an uncoordinated dysregulation of the ECM in senescent tenocytes, rather than coordinated activation or suppression of ECM-producing pathways, with fold-changes of ECM gene expression significantly more spread than the transcriptome as a whole, as confirmed by Browne-Forsythe test for equality of variances (Figure 3D, p < 0.0001).

Given these high-magnitude and significant changes, and that the key function of tenocytes is to maintain tendon ECM and its resilience to mechanical load, we focused our downstream analysis on ECM genes whose protein product has a known function in tendon (Screen et al., 2015). The interaction network in Figure 3E illustrates the data from tendon ECM genes (Table 1), where gene nodes are sized according to the expression level we detected in Lo-P cells, coloured according to the fold-change in Hi-P cells, and network edges weighted according to the StringDB interaction strength between protein products (Szklarczyk et al., 2023) (extended interaction network of the full matrisome is shown in Supplemental Figure 2B). Highlighted in the interaction network in Figure 3E are transcripts corresponding to collagens, fibril organisers, and elastic fibres; and the matrix metalloproteinases (MMPs) responsible for protein turnover of these functional groups are also included. High-magnitude changes in each of these functional groups demonstrates that Hi-P populations, which represent mixed populations of healthy and senescent cells, dysregulate tendon ECM synthesis, and likely dysregulate tendon ECM turnover.

**Table 1.** Expression of genes whose protein products have known function in tendon extracellular matrix (Screen et al., 2015).

| Gene | Expression level | Log<sub>2</sub> fold-change (Lo-P → Hi-P) | FDR-adjusted p value |
| --- | --- | --- | --- |
| COL1A1 | 596455.19 | -0.05 | 0.94 |
| FN1 | 302188.25 | 0.61 | 0.32 |
| COL1A2 | 301967.27 | 0.16 | 0.77 |
| THBS1 | 224873.39 | 0.47 | 0.61 |
| COL3A1 | 106355.76 | 0.22 | 0.74 |
| COL5A1 | 8915.00 | 0.32 | 0.61 |
| COL5A2 | 44977.53 | 0.51 | 0.31 |
| COL5A3 | 27.48 | 1.34 | 0.27 |
| COL11A1 | 4707.95 | 1.28 | 0.12 |
| COL11A2 | not expressed |  |  |
| **COL14A1** | **1213.29** | **-4.17** | **0.00028** |
| **DCN** | **28477.97** | **-4.02** | **1.6E-08** |
| PRG4 | 27.90 | 0.75 | 0.76 |
| **TNC** | **3362.20** | **3.50** | **0.0023** |
| **COMP** | **20741.87** | **-6.50** | **2.3E-06** |
| TNMD | not expressed |  |  |
| BGN | 35739.64 | 0.12 | 0.89 |
| FMOD | 2286.47 | -0.64 | 0.84 |
| LUM | 30848.63 | -0.35 | 0.52 |
| **VCAN** | **11169.42** | **1.06** | **4.9E-08** |
| ACAN | 1332.56 | -4.79 | 0.14 |
| **ELN** | **447.16** | **-5.10** | **1.1E-08** |
| FBN1 | 55118.59 | 0.06 | 0.90 |
| **FBN2** | **69.38** | **2.34** | **0.026** |

### Senescence compromises tenocyte wound healing, cell-matrix interactions, and cell-cell signalling

Next, we sought to understand the tendon tissue health implications of these senescence- associated changes in tenocytes, performing several assays of typical tenocyte function. We began with a simple scratch assay, wherein an artificial wound is introduced to confluent *in vitro* cultures, with proliferation inhibited through serum restriction. We found that Hi-P cells were far less efficient at closing the artificial wound site, closing 2.50 % of the wound area per hour on average as opposed to 5.66 % in Lo-P cells (Figure 4A, Supplemental Videos 1-2, p = 0.0126). Lo-P cells fully restored the culture monolayer by 48 hours, with Hi-P cells still yet to fully close the wound site after 72 hours (Supplemental Figure 3A).

**Figure 4.**
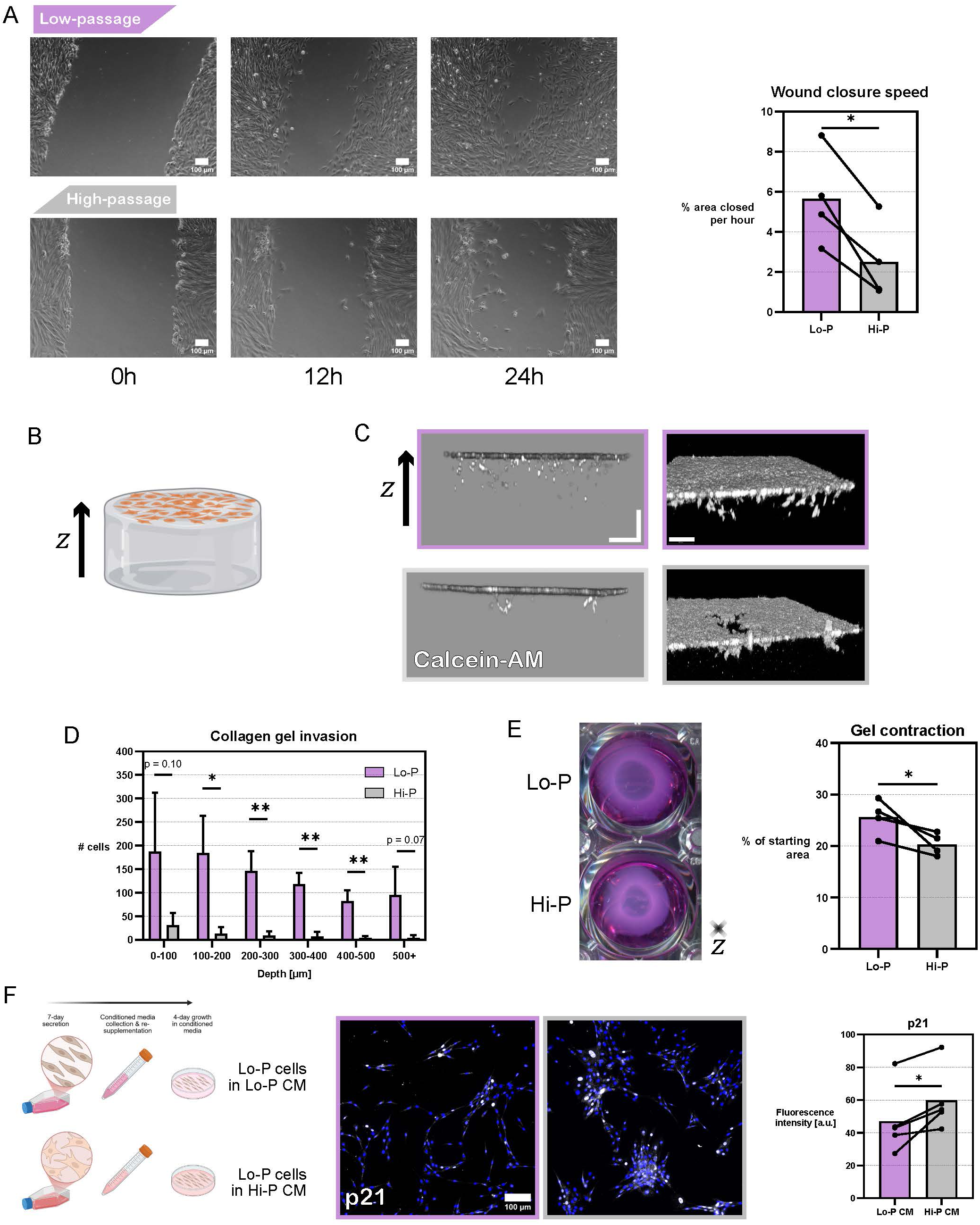
Senescence compromises tenocyte function. **(A)** Scratch assay showing live microscopy images of low- (Lo-P, purple) and high- (Hi-P, grey) passage cultures in the 24 hours following introduction of an artificial would site. N = 4 **(B)** Schematic, Lo-P and Hi-P tenocytes are seeded on top of a collagen gel, with gel invasion and contraction measured after one week. **(C)** Confocal microscopy images with live cell marker calcein-AM demarcating cell invasion into collagen gels after one week of culture. Left: orthographic projection showing collagen gel invasion. Right: isometric projection showing collagen gel surface integrity. **(D)** Quantification of the distance cells invade into the collagen gel after one week. Note only invading cells are quantified, i.e. a depth of 0 µm begins just below the gel surface. Data are represented as mean ± SD. N = 4. **(E)** Top-down view and measurement of the contraction of collagen gels after one week, quantified as a percentage of the initial area. N = 4. **(F)** Left: schematic, conditioned media (CM) is collected from Lo-P and Hi-P tenocytes after one week of culture before being re-supplemented and used as growth media for Lo-P cultures. Right: immunofluorescence of p21 protein in Lo-P tenocytes treated for four days with either Lo-P (purple) or Hi-P (grey) conditioned media. Arbitrary units. N = 5. Unless otherwise specified, bars represent means of biological replicates, and p values show the results of a paired two-tailed t test (∗ < 0.05, ∗∗ < 0.01).

Given the highly collagenous nature of tendon tissue, and senescence-associated changes in MMP expression seen through our RNA sequencing data, we investigated how Lo-P and Hi-P tenocytes interact with a collagen matrix. Seeding equal numbers of cells on top of a collagen gel (Figure 4B, proliferation inhibited through serum restriction), we first observed how efficiently cells are able to migrate through the collagen matrix over the course of one week. As seen in 2D scratch assays, Hi- P tenocytes were also much less able to migrate through the 3D collagen gel (Figure 4C-D). Further, there appeared to be differences in the mode of action of movement into the gel. Figure 4C demonstrates that while Lo-P tenocytes migrate through the gel as single cells, largely leaving the gel surface intact; Hi-P tenocytes move into the gel in clusters, leaving craters at the site of migration, suggesting a degradation of the collagen gel rather than migration through it. Indeed, this observation agrees with sequencing data showing markedly increased synthesis of collagen-specific MMPs (Figure 3E).

In a separate assay, with cells again seeded on top of collagen gels, gels were separated from the edges on the culture vessel and gel contraction by tenocytes was analysed over one week. Despite activation of inflammatory pathways in Hi-P tenocytes (Figure 3A), Hi-P tenocytes were significantly more contractile than Lo-P, contracting to 20.3 % of the original gel area versus 25.6 % (Figure 4E, p = 0.0495), which was associated with an upregulation of IL-1 and IL-6 family proteins (Supplemental Figure 3B). Interestingly, IL-6 has been shown to induce actomyosin expression (Gallucci et al., 2006), and our RNA sequencing data revealed consistent, if moderate, increases of actomyosin components in Hi-P tenocytes (Supplemental Figure 3C). Therefore, we conclude that increased collagen matrix contraction in Hi-P populations is likely due to higher contractility in senescent tenocytes.

Figure 1C suggests that even in aged tendon, the proportion of senescent tenocytes is relatively low (∼4 %). However, previously it has been shown that the proinflammatory, ECM-modifying SASP can negatively affect, and even induce senescence in, neighbouring healthy cells within days (Acosta et al., 2013). To investigate the effects of the SASP in tenocytes, seven-day conditioned media was collected from Lo-P and Hi-P tenocyte cultures before administrating each to Lo-P tenocytes. Indeed, after just four days of exposure to Hi-P conditioned media, Lo-P tenocytes exhibited significantly increased p21 expression versus those treated with Lo-P conditioned media (Figure 4F, p = 0.0229), as well as a minor increase in β-galactosidase staining (Supplemental Figure 3D). This is an important result, as it demonstrates that even with adequate clearance machineries leading to transient senescent populations *in vivo*, these cells will quickly have a deteriorative effect on the tissue.

### Extracellular matrix remodelling increases in senescent tenocytes

To validate our findings that Hi-P tenocytes dysregulate the synthesis of ECM proteins and upregulate MMPs at the gene level, we performed stable isotope labelling of amino acids in cell culture (SILAC) and harvested the decellularised matrix deposited by Lo-P and Hi-P cultures. In brief, cells were cultured in “light” media with standard formulation for one week, before being swapped for “heavy” media of the same formulation except for lysine, which was swapped for the stable ^13^C isotope (Figure 5A). At time points of 0, 48, and 72 hours of heavy media incubation, tissue culture vessels were decellularised using detergents, and deposited ECM was collected through mechanical scraping. Immunofluorescence of collagen in decellularised culture vessels confirmed the preservation of deposited ECM (Figure 5B).

**Figure 5.**
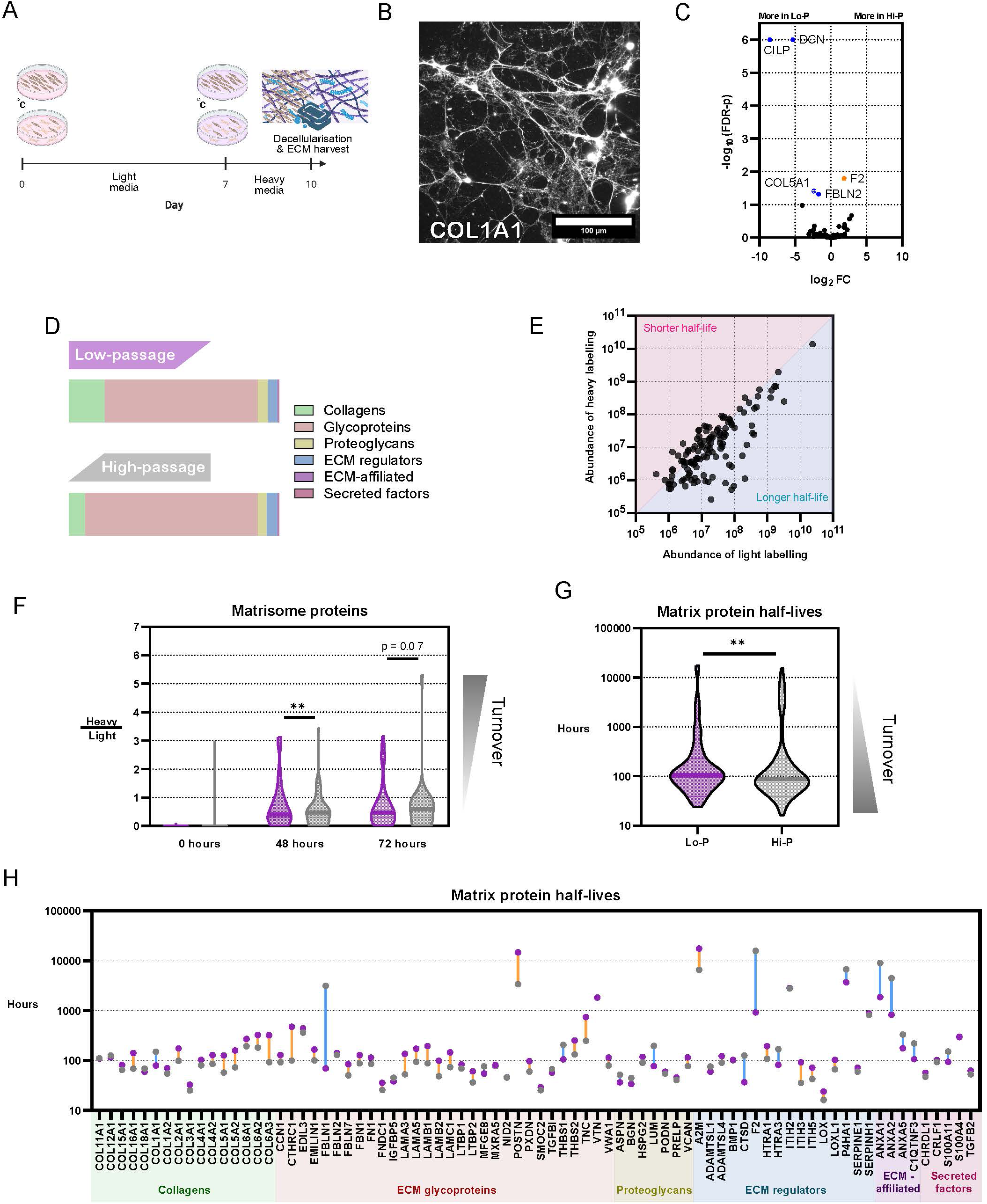
Proteomic changes to extracellular matrix composition and turnover in senescent tenocytes. **(A)** Schematic. Low- (Lo-P, top) and high-passage (Hi-P, bottom) tenocytes were incubated on tissue culture plastic (TCP) for one week in media with standard “light” lysine. Media was then swapped for the “heavy” media containing stable isotope ^13^C-lysine for up to 72 hours. TCP vessels were decellularised at time points of 0h, 48h or 72h after heavy isotope labelling and deposited matrix was collected for proteomic analysis. **(B)** Immunofluorescence of TCP-bound type I collagen stained after decellularisation. **(C)** Volcano plot of proteomic changes in deposited matrix between Lo-P and Hi-P cells. Coloured points satisfy a false discovery rate-corrected p value < 0.05. Blue indicates significant downregulation in Hi-P, orange indicates significant upregulation in Hi-P, p values from mass spectrometry proteomics are calculated as described in Methods. N = 4. **(D)** The protein composition of deposited matrix from Lo-P and Hi-P cells. N = 4. **(E)** The abundances of light- and heavy labelled proteins in Lo-P cells after 72 hours of heavy labelling. **(F)** Violin plot of heavy-to-light ratios of matrisome (Shao et al., 2023) proteins after 0, 48, and 72 hours of heavy labelling, deposited by Lo-P (purple) and Hi-P (grey) cells. N = 4. **(G)** Violin plot showing the calculated half-lives of matrisome proteins. N = 4. **(H)** The half-lives of individual deposited matrisome proteins by Lo-P (purple) and Hi-P (grey) cells. Blue lines indicate proteins with increased half-life in Hi-P cultures, orange lines indicate proteins with decreased half-life in Hi-P cultures. Unless otherwise specified, bars represent sample medians, and p values show the results of a Wilcoxon signed rank t test (∗∗ < 0.01).

Matrisome proteins (Shao et al., 2023) which were detected in at least three of four biological replicates were quantified, with 5 % of 101 proteins significantly regulated between Lo-P and Hi-P deposited ECM after adjusting for false discovery. Decorin, cartilage intermediate layer protein (CILP), type V collagen, and fibrillin-2 were significantly underrepresented in Hi-P deposited matrix, while prothrombin was significantly overrepresented in Hi-P deposited matrix (Figure 5C). The highest magnitude protein changes, decorin and CILP, followed the pattern seen in transcriptional changes (Supplemental Figure 2B). Changes were also seen in the overall composition of deposited ECM, with collagens making up 17 % of Lo-P ECM but just 8 % of Hi-P ECM, whereas glycoproteins made up 73 % of Lo-P ECM but rose to 82 % in Hi-P ECM (Figure 5D, Table 2).

**Table 2.** The composition of deposited matrix proteins from Lo-P and Hi-P tenocytes, according to matrisome division (Shao et al., 2023).

| Matrisome division | Lo-P % | Hi-P % |
| --- | --- | --- |
| Collagens | 17.08 | 7.86 |
| Glycoproteins | 72.91 | 82.16 |
| Proteoglycans | 4.79 | 4.12 |
| ECM regulators | 4.60 | 5.29 |
| ECM-affiliated | 0.43 | 0.35 |
| Secreted factors | 0.20 | 0.22 |

Through SILAC, protein half-lives were calculated by comparing the abundances of heavy- and light-labelled proteins across time points, as in previous work (Figure 5E) (Pickard et al., 2025; Schwanhäusser et al., 2009). As proof of principle, the 0-hour time point demonstrated an absence of detected heavy labelling, and heavy-to-light protein ratios increased with time (Figure 5F). At both 48-hour and 72-hour time points, heavy-to-light ratios were increased in Hi-P versus Lo-P (48-hour median ratio: Lo-P 0.4071, Hi-P 0.4733, p = 0.0036; 72-hour median ratio: Lo-P 0.4728, Hi-P 0.5874, p = 0.0698). 77 matrix proteins met the inclusion criteria for half-life analysis. Calculated protein half-lives were decreased overall in Hi-P deposited ECM, indicating an increased rate of ECM remodelling (Lo-P median half-life 88 hours, Hi-P median half-life 107 hours, p = 0.0049, Figure 5G). Individually, 52 proteins had shorter half-lives in Hi-P versus Lo-P compared to 25 with longer half-lives (Figure 5H).

### Senotherapeutic treatment of high-passage tenocytes alleviates phenotypes of senescence

Finally, having thoroughly characterised the expressional and behavioural changes undergone by senescent tenocytes, we set out to trial a number of senotherapeutic compounds to alleviate the senescent phenotype. We selected Q+D as examples of senolytics, which remove senescent cells, dRSV as a senoreverter, which restores proliferative capacity, and Adz as a senomorphic, which suppresses the SASP.

Toxicity assays were performed by administering varying concentrations from 10 nM to 100 µM of each compound to Lo-P cultures, three times over the course of seven days (Supplemental Figure 4A), in reduced serum (0.5 %). Working concentrations for further senotherapeutic analysis were chosen as the strongest concentrations which showed no adverse effects on culture health (as observed from bright field microscopy), cell viability (as quantified using Trypan blue), or cell metabolism (as quantified from resazurin metabolism). Working concentrations were selected as follows: Q, 100 nM; D, 5 nM (adverse effects seen at lowest dose tested); dRSV, 100 µM (no adverse effects at any dosage tested); Adz, 1 µM. While we opted for a seven-day treatment to allow beneficial effects to manifest, shorter and stronger doses of Q+D cocktail were also trialled. 24-hour administrations of several Q+D cocktails in respective concentrations of 0.01 – 1 mM and 0.1 – 10 µM however were all found to be lethal to Lo-P tenocyte populations (Supplemental Figure 4B).

Hi-P populations were subjected to the same programme as was used for toxicity assays, namely three treatments over the course of seven days in reduced serum (0.5%). All cultures were then switched to standard culture medium (10 % serum) for three further days such that any beneficial effects on cell proliferation would become apparent. After this time, treated cells were compared to Lo-P and Hi-P vehicle-only controls.

Beneficial effects on cell proliferation were measured through quantification of Ki-67-positive cells and numbers of viable cells (Figure 6A-C). Significant effects on cell proliferation were only seen in the comparison of Ki-67-positive cells between Q+D treated cultures and vehicle-only controls (p = 0.0304), however nonsignificant positive effects on numbers of Ki-67-positive cells and numbers of viable cells were seen with each senotherapeutic compound tested. β-galactosidase staining revealed a decrease in the numbers of senescent cells in dRSV-treated cultures, versus no apparent change in cultures treated with Q+D or Adz (Figure 6D, supplemental replicate images in Supplemental Figure 4C), however no change in p21 protein expression was detected for any of the three treatments (Supplemental Figure 4D).

**Figure 6.**
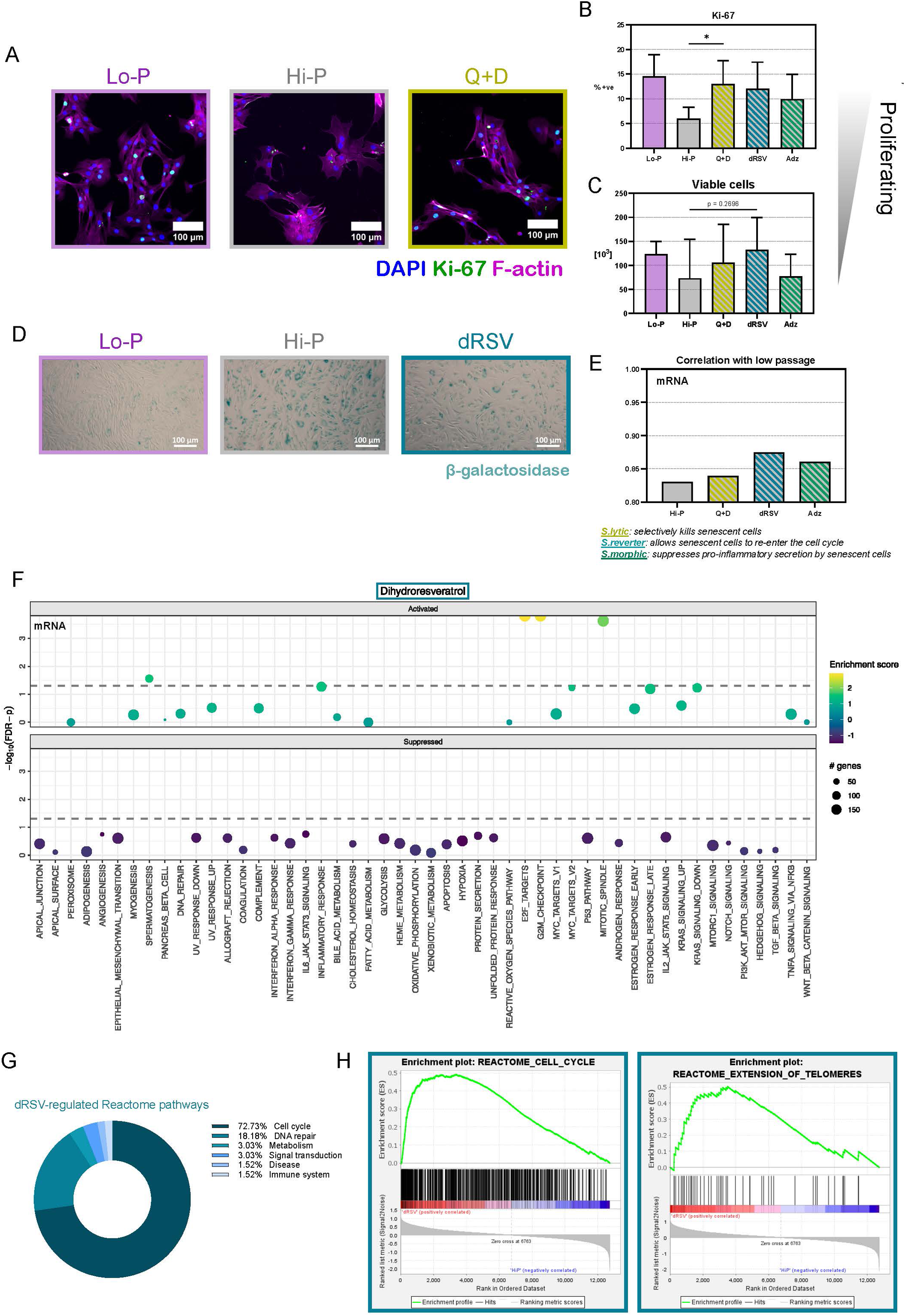
Senotherapeutic treatment has regenerative effects on high passage tenocytes. **(A)** Immunofluorescence of S-phase marker Ki-67 protein in tenocytes isolated from either low- (vehicle only, Lo-P), high-passage (vehicle only, Hi-P), or high-passage (treated with quercetin + dasatinib, Q+D) tenocytes. **(B,C)** Quantification of Ki-67 immunofluorescence (B) and viable cell number (C, starting cell number: 19,000) after seven-day treatment + three-day recovery in Hi-P cells treated with senolytics (Q+D), senoreverter (dRSV), or senomorphic (Adz). (B) N = 5, (C) N = 6. Data are represented as mean ± SD, and p values show the results of a paired two-tailed t test (∗ < 0.05). **(D)** Bright field images of cells stained for senescence-associated β-galactosidase after seven-day treatment + three-day recovery in Lo-P and Hi-P tenocytes, and Hi-P tenocytes treated with senoreverter (dRSV). N = 5, see supplementary replicate images in Supplemental Figure 4. **(E)** Pearson correlation coefficient comparing the transcriptome of low-passage tenocytes with the transcriptomes of high-passage cells either untreated or treated with senolytics (Q+D), senoreverter (dRSV), or senomorphic (Adz). N = 3. **(F)** Gene Set Enrichment Analysis (GSEA) of enrichment in the Hallmark gene sets in high-passage cells treated with the senoreverter dihydroresveratrol versus vehicle-only high-passage cells. Dashed line indicates a false detection rate-corrected p value < 0.05 (Subramanian et al., 2005). N = 3. **(G)** GSEA enriched Reactome pathways in high-passage cells treated with the senoreverter dihydroresveratrol (dRSV) versus vehicle-only control. N = 3. **(H)** GSEA enrichment of the Reactome pathways Cell cycle and Extension of telomeres versus vehicle-only control. N=3.

We also performed RNA sequencing to better characterise the effects of each senotherapeutic. By calculating Pearson correlation coefficients, we found that treatment with each senotherapeutic compound improved the transcriptional similarity between Hi-P and Lo-P cells versus vehicle-only control (Figure 6E), while also reducing the number of differentially expressed genes compared to Lo-P cells (Supplemental Figure 4E). dRSV was the most effective senotherapeutic by these metrics, with the transcriptome of dRSV-treated Hi-P tenocytes correlating most closely with Lo-P tenocytes (Pearson’s *r* = 0.87 versus 0.83 in vehicle-only control), and having the fewest significantly regulated genes versus Lo-P tenocytes (1,665 genes versus 2,012 in vehicle-only control). Through GSEA we identified significant changes in dRSV-treated Hi-P populations only, with four Hallmark gene sets activated (Liberzon et al., 2015) (Figure 6F, Supplemental Figure 4F-G). Three dRSV- activated gene sets related to proliferation: E2F targets, G2/M checkpoint, and Mitotic spindle assembly; and crucially two of these were identified as suppressed pathways in Hi-P populations compared to Lo-P (Figure 3A). A fourth gene set, Spermatogenesis was significantly activated, however among the top 15 enriching genes were several relating to cell cycle progression, including CDK1, BUB1, CCNB2, CCNA1, and RFC4, and so we consider this a misnomer in this context. We therefore also compared dRSV data against Reactome pathways, which are hierarchically structured and much more specific than Hallmark gene sets, at the cost of redundancy between pathways (Ragueneau et al., 2026). 66 Reactome pathways were significantly regulated after FDR correction in dRSV-treated Hi-P populations, mostly consisting of pathways within the cell cycle (48 pathways, 73 %) and DNA repair (12 pathways, 18 %) hierarchies (Figure 6G). Indeed, the top-tier pathway Cell cycle was strongly activated (FDR-corrected p < 0.0001), as well as Extension of telomeres (FDR-corrected p = 0.0252), which corroborates previous reports of dRSV acting to increase telomere length (Latorre et al., 2017) (Figure 6H).

## Discussion

In this study, we showed that tenocytes isolated from aged tendons express multiple markers of senescence, and established a replicative senescence model to determine functional consequences of tenocyte senescence. Using this model, we reported that senescent tenocytes have a similar profile to senescent cells observed in other tissues, such as dermal fibroblasts (Nan et al., 2025), and are proinflammatory and contractile, with impaired wound healing capacity. Additionally, these cells dysregulate tendon matrix synthesis and turnover, and secrete factors which induce senescence in healthy tenocytes. By characterising an array of senotherapeutics, we showed that each compound assessed improved the transcriptomic similarity between senescent and healthy cells, with dRSV in particular restoring pathways associated with proliferation at the level of transcription and reducing markers of senescence.

Previously we have reported evidence of senescent tenocytes accumulating in aged tendon (Zamboulis et al., 2024), and here we have shown that tenocytes isolated from aged tendons also exhibit markers of senescence *in vitro*. A reduced growth rate in tenocyte populations isolated from aged tendons could indicate either a decreasing proliferative fraction or an increasing cell cycle length with age, however concomitant results of increased p21 and β-galactosidase expression suggest the former of these indications. By inducing senescence through serial passaging in tenocytes isolated from young tendons, we have replicated these senescence phenotypes in Hi-P cells, yielding an *in vitro* model to directly study the consequences of age-associated tenocyte senescence.

### Senescent tenocytes are proinflammatory and dysregulate extracellular matrix remodelling and cell-matrix interactions

Here we have demonstrated that senescent tenocytes are proinflammatory and upregulate TNF-α signalling pathways, underpinned by upregulation of members of the proinflammatory IL-1 and IL-6 families (Al-Qahtani et al., 2024; Kaneko et al., 2019).

While senescent tenocytes had a proinflammatory phenotype, they also had increased contractile ability. Studies in smooth muscle cells have linked inflammation to increased cell contractility (Khalfaoui and Pabelick, 2023; Lamb et al., 2024). IL-6 in particular has been shown to induce actomyosin machinery (Gallucci et al., 2006) and we observed broad upregulation of actomyosin components at the gene level, including significant upregulation of the contractile proteins myosin- 10 and myosin-14 (Chinthalapudi and Heissler, 2024; Murrell et al., 2015). While the senescent cell population in adult tendon is small, together these results suggest senescent tenocytes contribute to driving low-grade inflammation and increasing local tissue stiffness and contraction. This increased contractility in Hi-P cells likely combines with characteristic large focal adhesions in senescent cells (Brauer et al., 2023; Shin et al., 2020) to produce the impaired 2D and 3D motility seen in Figures 4A-D, with possible relevance to evidence of poorer tendon healing in aged patients (Ducasse and Collin, 2024; Rashid et al., 2017), with age being the strongest predictor of outcome after Achilles tendon rupture (Carmont et al., 2020).

Figure 3C-E shows evidence of significant dysregulation of tendon ECM synthesis by Hi-P cells, as well as hallmark upregulation of MMPs (Hernandez-Segura et al., 2018), which has also been observed in cyclically loaded tendon explants (Dudhia et al., 2007). In particular, Hi-P tenocytes experience marked upregulation of tenascin C, versican, and fibrillin 2; and marked downregulation of type XIV collagen, decorin, cartilage oligomeric matrix protein (COMP), and elastin. Decorin’s glycosaminoglycan (GAG) chains bind to collagen I fibrils and set the distance between fibrils; shortening of these chains is associated with structural changes in aged skin (Li et al., 2013; Nomura, 2006) and tendon (Dunkman et al., 2013). COMP is also thought to be a molecular chaperone for collagen fibrillogenesis (Halász et al., 2007) and previous evidence has shown COMP decline both in senescent cells and in ageing tendon tissue (Smith et al., 1997). Interestingly, transduction of COMP into senescent murine vascular smooth muscle cells reduced levels of p53, p21, and p16 (Wang et al., 2016). Downregulation of decorin has been observed in senescent fibroblasts in other tissues (Mavrogonatou et al., 2021), and previously we have shown that elastin abundance and organisation are both decreased in aged tendons (Godinho et al., 2017). Further, loss of synthesis of type XIV collagen and elastin are compounded by significantly increased synthesis of their turnover machineries MMP1 and MMP12 (log_2_ fold-changes 9.95, 11.1 and FDR-adjusted p values 7.12×10^-5^, 5.12×10^-3^ respectively) (Hey and Linder, 2024).

Half-lives of tendon ECM proteins are generally considered to be extremely long. Our previous (Thorpe et al., 2010) measurement of aspartic acid racemisation in the SDFT suggested collagenous proteins have a half-life of approximately 200 years in tendon, while the value for non-collagenous proteins is approximately 2 years. A study utilising the temporal dynamics of atmospheric ^14^C levels across the last century (Heinemeier et al., 2013) also found extremely low levels of ECM remodelling of the Achilles tendon core in adulthood, also reporting high biological variability in tendon turnover rates consistent with injury-induced ECM remodelling. However in a thermally dynamic tissue such as tendon these values may be overestimates and still allows for faster turn-over of a small, but structurally more important, component of the matrix, such as the interfascicular matrix, as previously proposed in mouse tendon (Chang et al., 2020).

In agreement with these studies, we found most of the ECM deposited by cells consists of glycoproteins rather than collagen (Figure 5D), despite the majority of in vivo matrix being made up by collagens (65-80 %), adding further evidence that tendon ECM remodelling primarily revolves around non-collagenous proteins. The role of glycoproteins in age-associated tendon degeneration has yet to be established (Kostrominova and Brooks, 2013; Kwan et al., 2023), however here we present evidence that senescent tenocytes increase the deposition of glycoproteins at the expense of collagen deposition. Reduced collagen production by senescent cells has been observed in human skin fibroblasts (Brauer et al., 2023) and murine tenocytes (Stowe et al., 2024), but interestingly we observed this senescence associated reduction in collagen deposition independently from any substantial change in collagen transcription (Figure 3E).

Increased synthesis of MMPs in Hi-P tenocytes was associated with higher rates of ECM protein turnover by Hi-P cells (Figure 5F-H), and resulted in crater formation within collagen gels (Figure 4C). These results may be indicative of a degenerative effect of senescent cells on ageing tendon. The combined increase in collagen turnover and decrease in collagen deposition by Hi-P cells may have crucial implications for the tendon core in ageing tendon, given the lack of regeneration in this part of the tissue (Heinemeier et al., 2013; Thorpe et al., 2010). Indeed, aged tendons have been shown to accumulate damaged, partially cleaved collagen (Thorpe et al., 2010), and cyclical loading of aged tendon explants results in increased tendon matrix disruption compared to younger tendon (Dudhia et al., 2007). As well as these changes in ECM deposition and remodelling, Hi-P tenocytes secreted factors which induced senescence in healthy tenocytes within days (Figure 4F), replicating previous results in human fibroblasts (Acosta et al., 2013).

While we and others have shown that aspects of the senescent fibroblast phenotype are conserved across induction methods (Hernandez-Segura et al., 2017) (Supplemental Figure 1D), questions remain about the mechanism by which senescence is induced in ageing tendon. Studies of cell proliferation in adult tendon are scarce, however in adult ovine calcaneal tendon it has been shown that approximately 5 % of tenocytes are Ki-67-positive at any given time (Russo et al., 2015). While further study in this area is needed, this would suggest a dynamic *in vivo* tenocyte population with several doublings per year, and in Supplemental Figure 1B we demonstrate that tenocyte populations isolated from young adult tendons are capable of at least 20 doublings, and likely around 50 doublings.

Further, while the scope of this work has been a mixed population of tenocytes, we have also previously shown evidence of senescence in tendon microvascular cells and interfascicular matrix cells (Zamboulis et al., 2024). Indeed, we have characterised age-associated changes to tendon microvasculature (Iwasaki et al., 2026) and future work should seek to investigate senescence in tendon microvascular cells. However even on optimised substrates, *in vitro* expansion potential of interfascicular matrix fibroblasts is extremely limited, preventing a parallel study in these cells, and remains a challenge to be overcome (Grossemy et al., 2026).

### Senotherapeutic treatment restores healthy cell phenotypes in senescent tenocytes

Treatment options for chronic tendon injury are limited in their effectiveness (Nourissat et al., 2015; Smith, 2024). The benefits of platelet-rich plasma therapy are contentious, and methods including corticosteroids, thermocauterisation, shockwave, electrostimulation, ultrasound, and laser lack evidence of efficacy while also causing pain and other potentially damaging effects.

More effective treatment strategies revolve around resolving inflammation and promoting optimal ECM remodelling and organisation of scar tissue. These include cold and/or compression to resolve inflammation in acute stage, while in the subacute and chronic stages, there is increasing clinical evidence that administration of mesenchymal stem cells can reduce rates of reinjury (Godwin et al., 2012; Salz et al., 2023), thought to be via these cells’ immunomodulatory capacity. Previously, we have shown that senescence in these cells adversely affects their immunomodulatory phenotype (Llewellyn et al., 2023), and that both tenocyte and MSC behaviours are influenced by the structure and composition of their surrounding matrix (Clements et al., 2016).

Here we present the results of three distinct senotherapeutic compounds targeting senescent tenocytes in Hi-P cell populations. Treatment time courses were set out to demonstrate differences in the mechanisms of action of these compounds. Previous characterisations of our three candidate compounds suggested that only the senolytic cocktail of quercetin and dasatinib would induce cell death selectively in senescent cells (Hawthorne et al., 2024; Takaya and Kishi, 2024; Xu et al., 2018), and by carrying out an initial seven-day treatment in reduced serum, we expected to see a reduction in cell numbers following treatment of Q+D only. However, substantial cell death was not seen in Hi-P tenocytes treated with any of the senotherapeutic compounds trialled (Figure 6C, Supplemental Figure 4C). Further, by testing a range of concentrations, we were not able to demonstrate any selective senolytic effect of Q+D in tenocytes (Supplemental Figure 4B). Q or Q+D have also been shown previously to decrease senescence markers in aged tendon (Cai et al., 2025), suppress inflammation and SASP secretion in tendon injury (Wang et al., 2026), and enhance tendon healing (Yurteri et al., 2025); although in a study of patients with chronic tendinopathy, tendons with elevated markers of senescence were unaffected by exposure to Q+D (Bühler et al., 2022). At the conclusion of a ten-day treatment, consisting of seven days’ treatment in reduced serum and three days of growth, we observed Q+D significantly increased the proliferation of Hi-P tenocytes (Figure 6A-B) but had no noticeable effect on β-galactosidase expression (Supplemental Figure 4C). This ten-day time course at the highest concentration sublethal to Lo-P tenocytes (Supplemental Figure 4A) was also insufficient to induce any pathway-level transcriptional changes in Hi-P tenocytes (Supplemental Figure 4F). Adz-treated Hi-P tenocytes similarly showed no significant regulation of pathways (Supplemental Figure 4G) despite a minor improvement in cells’ transcriptional resemblance of Lo-P tenocytes (Figure 6E, Supplemental Figure 4E), as well as no significant effect on cell proliferation and viability (Figure 6B-C) or β-galactosidase and p21 expression (Supplemental Figure 4C-D) versus vehicle-only control. It is important to note that, to the authors’ knowledge, no previous studies have investigated the therapeutic effect of Q+D or Adz in cells of equine origin, therefore there may be species differences that limit the effectiveness of these compounds in the horse.

dRSV proved to be the most broadly beneficial senotherapeutic of the compounds trialled. Treatment of Hi-P tenocytes resulted in a clear reduction in β-galactosidase expression (Figure 6D, Supplemental Figure 4C) and modest improvements in cell proliferation and viability (Figure 6B-C). Through RNA sequencing of dRSV-treated Hi-P tenocytes we also found that the transcriptomes of these cells more closely resembled Lo-P tenocytes (Figure 6E, Supplemental Figure 4E) and saw significant upregulation of cell cycle pathways versus vehicle-only control (Figure 6F), in line with our previous results in equine MSCs (Tamura et al., 2023) and corticosteroid-induced senescent equine tenocytes (Heidari et al., 2024). Deeper analysis using Reactome pathways corroborated previous evidence (Da Luz et al., 2012; Latorre et al., 2017) that the mechanism of these actions is via extension of telomeres (Figure 6H). It should be noted that even modest influences may be sufficient to slow tendon degeneration sufficiently to act as an effective prevention of tendinopathy.

## Conclusion

Tendon injuries pose significant challenges to both human and veterinary medicine, with risk of injury increasing with age and injured tendons remaining recalcitrant to treatment. Taken together, these results highlight a degradation cycle of age-associated senescent tenocytes. Via inflammation signalling, ECM contraction, and ECM degradation, senescent tenocytes produce a signal similar to that seen in tendinopathy, and may drive phenotypes in ageing associated with high likelihood of tendon injury. Across a panel of senotherapeutic compounds, seven-day treatment with dRSV had a beneficial effect on Hi-P tenocytes across several measures. In an ageing population, these findings have important implications for the prevention and treatment of tendon injury.

## Methods

### Resource availability

#### Lead contact

Further information and requests should be directed to and will be fulfilled by the lead contact, Dr. Chavaunne Thorpe

### Materials availability

This study did not generate new unique reagents.

### Data and code availability

Data: All raw data reported in this paper is available at online repositories via the accession numbers herein. All processed data will be shared by the lead contact upon request.

Code: This paper does not report original code.

Any additional information required to reanalyse the data reported in this paper is available from the lead contact upon request.

### Experimental model and study participant details

Forelimbs, distal to the carpus, were collected from horses euthanised for reasons unrelated to this project from a commercial abattoir. Sample collection was approved by the Royal Veterinary College’s Clinical Research Ethical Review Board (URN 2021 2077-2). ∼5cm pieces of SDFT were harvested from the mid-metacarpal region of the forelimb within 24 hours of euthanasia. Tenocytes were isolated from the SDFTs of either young, skeletally mature horses (3-6 years) or aged horses (18+ years).

## Method details

### Tendon digestion and primary tendon cell culture

SDFT pieces were washed several times in phosphate-buffered saline (PBS, Fisher Scientific #BP2944), the epitenon was removed, and the tendon core was finely minced (a few mm^3^) under sterile conditions. Samples were subjected to a two-step digestion in DMEM consisting of 1 mg/ml pronase E (Sigma-Aldrich #1074330005) for 1 h under agitation at 37 °C, followed by 0.5 mg/ml collagenase type 4 (Worthington #LS004188) and 1 mg/ml dispase II (Sigma-Aldrich #D4693) for 24 h under agitation at 37 °C. DMEM was supplemented with 100 µg/ml of the primary cell microbial agent Primocin (InvivoGen #ant-pm-05) during 24 h digestion and the first few days of culture.

Following digestion, samples were strained (70 μm filters) and centrifuged (400g, 5 min) to remove debris prior to seeding. Primary tenocytes were maintained on tissue culture-treated plastic, coated with 2-5 µg/cm² type I collagen (Corning #CLS354236) in low glucose DMEM (Gibco #31885) supplemented with either 10 % fetal bovine serum (Thermo Scientific #10500064) or 10 % donor horse serum (Life Science Production #S-004-US), 1 % Penicillin/Streptomycin cocktail (Gibco #11548876), and maintained in a humidified 37°C incubator with 5 % CO_2_. No changing of serum or serum batches occurred within individual experiments. Medium was changed thrice weekly, and cell cultures were dissociated using Accutase (Stem Cell Technologies #07920) and reseeded at 1,000 cells/cm² ad-hoc prior to confluency, approximately every 7 days.

All quantitative comparisons were made between cell populations from the same subject, matched Lo-P and Hi-P. Following digestion from tendon tissue, a portion of cells were kept in cryogenic storage (Lo-P) whilst subject-paired cells were passaged to a point of replicative senescence (Hi- P), designated according to changes in cell morphology and growth rate, and verified with a combination of markers such as β-galactosidase, metabolic activity, nuclear morphology, proliferation markers and cell cycle arrest markers. At this point Lo-P cells were reseeded into culture, allowing experiments to be performed on Lo-P and Hi-P paired populations concurrently. Cells from different subjects were never pooled and N numbers quoted refer to biological replicates (i.e. from different subjects). Details of individual subjects, and in which experiments they were used, are summarised in Table 3.

**Table 3.** Biological subject information.

| Subject ID | Age [years] | Experiments |
| --- | --- | --- |
| H2401 | 3 | Senotherapeutics, p21 ICC, scratch assays, gel assays, paracrine signalling assay, SILAC |
| H2402 | 4 | Gel assays, RNA sequencing (senescence), senotherapeutics, growth assays, p21 ICC, Ki-67 ICC, Lamin B1 ICC, metabolism assays, scratch assays |
| H2403 | 18 | Growth rate measurements, p21 ICC |
| H2405 | 6 | RNA sequencing (senescence), Senotherapeutics, growth rate measurements, p21 ICC, Ki-67 ICC, Lamin B1 ICC, metabolism assays, scratch assays |
| H2406 | 19 | Beta-galactosidase, growth assays |
| H2407 | 18 | Growth rate measurements, p21 ICC |
| H2409 | 4 | RNA sequencing (senescence), Growth rate measurements, p21 ICC, Ki-67 ICC, Lamin B1 ICC, metabolism assays, scratch assays |
| H2410 | 3 | RNA sequencing (senescence), Growth rate measurements, metabolism assays, Ki-67 ICC, Lamin B1 ICC |
| H2413 | 6 | Senotherapeutics, growth rate measurements, gel assays, paracrine signalling assay, SILAC |
| H2501 | 15 | p21 ICC |
| H2502 | 23 | p21 ICC |
| H2503 | 4 | Senotherapeutics, gel assays, paracrine signalling assay, SILAC |
| H2504 | 4 | Paracrine signalling assay |
| H2505 | 3 | Senotherapeutics, gel assays, paracrine signalling assay |
| H2506 | 2 | SILAC |

Unless otherwise stated, cells were maintained in low glucose DMEM (Gibco #31885049) supplemented with 10 % horse serum (Life Science Production #S-004-US) and 1 % penicillin/streptomycin (Fisher Scientific #11548876), in tissue culture-treated plastic (TCP) coated with ∼5 µg/cm² type I collagen (Sigma Aldrich #CLS354236-1EA), at 37 °C in 5 % CO_2_.

### Growth analysis

On each passaging, cells were reseeded at 1,000 cells/cm² and passaged prior to confluency. Cell number was determined via automated cell counting (DeNovix CellDrop) and the doubling time for the period since the previous passaging was calculated. Doubling time comparisons were serum batch-matched.

### Beta-galactosidase staining

Beta-galactosidase staining was carried out using the Beta galactosidase staining kit (Abcam, #ab65351) according to the manufacturer’s instructions.

### Immunocytochemistry

Cells were allowed a minimum of 48 hours to equilibrate to the culture substrate (#1.5 glass coverslips coated with collagen-I unless otherwise stated) following seeding prior to fixation in 4 % formaldehyde (Sigma-Aldrich #1.00496) for 10 min at room temperature. Following fixation, cells were washed twice with PBS. Cells were permeabilized in 0.3 % Triton-x100 (Sigma-Aldrich #T8787) in PBS for 20 min at room temperature and blocked with 3 % bovine serum albumin (Sigma-Aldrich #05482), 2 % horse serum, 2 % goat serum (Sigma-Aldrich #G9023) in PBS for 1 hour at room temperature. Cells were incubated with primary antibodies (Table 4) in PBS at 4 °C overnight, followed by five PBS washes. Cells were then incubated with 1 µg/ml DAPI (Abcam #ab285390) and appropriate secondary antibodies (1/500, Invitrogen) in blocking solution with Phalloidin-iFluor647 where applicable (Abcam #176759) for 1 hour at 37 °C. Cells were then washed five times using PBS, following which coverslips were rinsed in ultrapure water before mounting onto 1 mm thick glass slides using anti-fade mounting medium (Abcam #104135).

**Table 4.** List of primary antibodies used for immunocytochemical analyses.

| Antigen | Species | Company | Dilution |
| --- | --- | --- | --- |
| Ki-67, monoclonal | Rb | Thermo Scientific #MA5-14520 | 1/100 |
| p16, monoclonal | Rb | Cell Signalling Technology #D7C1M | 1/50 |
| p21, monoclonal | Rb | Abcam #ab109520 | 1/500 |
| p53, polyclonal | Rb | Fisher Scientific #BS-2090R | 1/50 |
| Collagen 1A1, polyclonal | Rb | Thermo Scientific #PA5-29569 | 1/100 |
| Lamin B1, polyclonal | Rb | Abcam #ab16048 | 1/500 |

### Microscopy

Bright field images were taken on a Zeiss Primovert microscope using either a Zeiss Axiocam 208 or a DinoEye Edge AM7025X camera. Immunofluorescence images were acquired with either a Nikon Eclipse Ni-E wide-field or a Leica SP8 confocal microscope. Live cell imaging was carried out using a Leica DMI6000 with a custom-built 37 °C, 5 % CO_2_ chamber. Cells were observed using a 20x objective over 72-hour time periods and with a 30 min interval between frames.

### Resazurin metabolism assays

Equal numbers of Lo-P and Hi-P cells were incubated with 200 µM resazurin (ApexBio #B6098) for two hours, following which the 570 nM absorbance was read using a Tecan Cyto-400 plate reader, indicative of the extent of metabolic conversion to resorufin. Results were normalised to resazurin administered to blank wells for an equal time.

### Scratch assays

Cells were seeded at 20,000 cells/cm² onto uncoated TCP. After being given 72 hours to form a stable monolayer, cultures were switched to reduced serum (0.5 %) to inhibit proliferation. A 20 µL pipette tip was scratched through the culture, followed by live microscopy for 72 hours. For each replicate, average measures of triplicate wells were used for downstream data analysis.

### Dexamethasone treatment

Cells were seeded onto collagen coated plates at 5,000 cells/cm² and incubated for 24 hours to allow cell attachment. Using established protocols (Martin et al., 2019), cells were then treated with 200 nM dexamethasone (Sigma-Aldrich #D4902) in 10 % serum media for (a) 72 hours for p16 immunolabelling, and (b) 60 hours for p21 and p53 immunolabelling. Following dexamethasone treatment, the media was replaced with fresh complete growth medium, and cells were cultured for an additional 24 hours.

### RNA isolation and sequencing

RNA was extracted from cell pellets using the RNeasy Mini kit (Qiagen #74134), as per the manufacturer’s instructions, and concentration was measured via spectrophotometer (Denovix CellDrop FX).

Library construction, quality control, sequencing, and statistical analysis were carried out by Novogene Co., Ltd. Messenger RNA was purified from total RNA using poly-T oligo-attached magnetic beads. After fragmentation, the first strand cDNA was synthesized using random hexamer primers followed by the second strand cDNA synthesis. The library was ready after end repair, A-tailing, adapter ligation, size selection, amplification, and purification. The libraries were constructed with an insert size of approximately 250–300 bp. After library quality control, different libraries were pooled based on the effective concentration and targeted data amount, then subjected to Illumina sequencing. Sequencing was performed on the NovaSeq X Plus platform using the PE150 (paired-end 150 bp) sequencing strategy. Raw data is available on the EMBL-EBI platform ArrayExpress with the accession numbers E-MTAB-17377 and E-MTAB-17394.

### Collagen gel assays

Collagen-I was diluted in low glucose DMEM to 2.5 µg/mL. The collagen solution was then placed in a chamber with a pool of ammonium hydroxide (Sigma-Aldrich #30501) for 3 min to adjust the pH and form a cured collagen gel. Gels were washed twice with PBS prior to cell seeding.

For invasion and contraction assays, 100,000 cells/cm² were seeded on top of collagen gels and maintained for one week in reduced serum (0.5 %). For collagen contraction assays, gels were detached from culture wells the day after seeding, using a 10 µL tip in a circular motion. Contracted gels were imaged after one week using an Epson Perfection V370 scanner, gel edges were demarcated by hand and compared to the well area.

Calcein-AM (Thermo Scientific #C1430) was used to demarcate live cells in collagen invasion assays as follows. Gels were rinsed twice with PBS followed by a 15-minute incubation with 1 µg/ml calcein-AM in DMEM, after which gels were again rinsed twice with PBS. Images were then taken with gels in PBS. For quantification of invasion depth, a z plane immediately below surficial cells was defined as the origin. 3D images were divided into 100 µm segments below the origin, and the numbers of Calcein-AM-positive cells in z-projections of each segment were counted using FIJI.

### Paracrine signalling assays

Lo-P and Hi-P cells were seeded at 50,000 cells/cm². The day after seeding, media was refreshed with a double volume of reduced serum (0.5 %) media and left for 7 days. Conditioned media was then collected, resupplemented to 10 % serum, and used as the medium for separate (subject-matched) cultures of Lo-P cells (seeded at 1,000 cells/cm²) for 4 days (with one refresh at day 2).

### In vitro stable isotope labelling and extracellular matrix isolation

SILAC DMEM Flex Media (Gibco #A2493901) was supplemented to match the formulation of low glucose DMEM, with the exception of lysine, where 100 mg/L of either light (Thermo Scientific #89987) or heavy (Thermo Scientific #89988) lysine was added as appropriate. SILAC media was also supplemented with 50 µg/L ascorbic acid (Sigma-Aldrich #A92902) to promote collagen secretion. SILAC media was also supplemented with 1 % Penicillin/Streptomycin cocktail and 0.5 % dialyzed foetal bovine serum (Gibco #A3382001).

At D0, 20,000 cells/cm² were seeded onto uncoated TCP in light media, refreshed at D1 and D4. At D7, light media was replaced with heavy media. On this day, light media-only controls were also harvested as described below. At either D9 or D10 (corresponding to 48 hour and 72 hour time points post-isotope labelling), working carefully throughout to not disturb deposited matrix, the culture dish was washed twice with PBS and once with ultrapure water, following which the culture vessel was incubated with 5ml 26.4 mM ammonium hydroxide (Sigma-Aldrich #30501) in PBS + 0.5 % Triton-x100 at 37 °C for 3 min. Dishes were rinsed thrice with PBS and then left in a fume cabinet for 30 min for residual liquid to evaporate. 300 µL 0.011 % sodium dodecyl sulphate (Sigma-Aldrich #L4390), 0.003 % sodium deoxycholate (Sigma-Aldrich #D6750) in 25 mM ammonium bicarbonate (Thermo Scientific #10207183) in ultrapure water was added to dishes, which were then scraped and the liquid collected.

### Mass spectrometry sample preparation and analysis

Proteins were extracted and digested from 100 µl of extracellular-matrix-enriched fractions using the standard micro S-Trap protocol, following the guidelines provided by ProtiFi. 100 mM Triethylammonium bicarbonate buffer (Sigma-Aldrich #T7408) containing sodium dodecyl sulfate (5% final concentration) was added to each sample at a 1:1 ratio. Proteins were reduced with dithiothreitol (Sigma-Aldrich #D0632), alkylated with iodoacetamide (Sigma-Aldrich #I1149), and acidified with phosphoric acid (Sigma-Aldrich #466123) to a final concentration of 2.5 % (as per manufacturer guidelines). Samples were then loaded onto S-Trap micro spin columns (ProtiFi #C002-MICRO) in binding buffer (ratio sample:buffer 1:6), where proteins were trapped and subsequently digested with 1 µg of trypsin (Sigma-Aldrich #T4799) at 37 °C overnight. Tryptic peptides were sequentially eluted, dried, and resuspended in 30 µl of 0.1 % formic acid (Sigma-Aldrich #106526) in LC-MS-grade water.

Peptides were analysed by LC-MS/MS using a Vanquish Neo UHPLC system (Thermo) coupled to an Orbitrap Ascend mass spectrometer (Thermo). The Vanquish Neo was operated in trap-and-elute mode using an Acclaim PepMap trap column (100 µm × 2 cm, ThermoFisher) and an EASY-SPRAY PepMap analytical column (50 cm × 75 µm, ES903, ThermoFisher). Peptides were trapped and separated using a 60-min gradient: 2–20 % solvent B (0.1 % formic acid in 100 % acetonitrile (Fisher #15329865)) over 45 min, followed by 20–35% solvent B over 15 min, at a flow rate of 300 nl/min.

Eluted peptides were analysed on the Orbitrap Ascend in data-dependent acquisition mode. MS1 spectra were acquired in the Orbitrap at 120k resolution over an m/z range of 380–1500, with an AGC target of 4e5 and an S-lens RF level of 30. The top 20 most intense precursor ions were selected, isolated in the quadrupole with a 2 m/z window, and fragmented by HCD at 30 % collision energy. MS/MS spectra were acquired in the Orbitrap at 15k resolution with a maximum injection time of 27 ms.

### Senotherapeutic treatments

For toxicity assays each compound was solubilised in DMSO and serially diluted in low glucose DMEM, while topping up DMSO to ensure equal vehicle concentration in all conditions and control. At D0, 1,000 cells/cm² at low-passage were seeded in normal culture media. Compounds were administered on D1, and refreshed on D3 and D6 before cells were imaged, incubated with resazurin, and tested for viability on D8. Working concentrations were chosen as the highest dosage for which no compromise was observed in visual cell health, cell metabolism, or cell viability.

To assess the senotherapeutic effects of each compound, at D0, 10,000 cells/cm² (low- and high-passage) were seeded in reduced serum (0.5 %) media. The course of treatment followed that of the toxicity assay, administering with either the identified dosages or DMSO controls. At D8, compound/vehicle was removed and cells were cultured in normal (10 % serum) media. At D11, cells were either fixed, counted, or harvested for sequencing.

### Quantification and statistical analysis Immunofluorescence quantification

Images for comparison were taken conserving the magnification, laser power, camera gain, and exposure time. 10 frames of view (FOV) were taken per sample, with FOV chosen by viewing DAPI signal only to reduce researcher bias. Nuclear segmentation was carried out using the FIJI plugin StarDist (Schmidt et al., 2018), subjected to a 50-500 µm² filter. The (background-subtracted) mean pixel intensity of secondary antibody signal within each nucleus was recorded, with the median of these values across all nuclei across all FOV taken to represent the sample for graphical and statistical purposes.

### RNA sequencing

Library construction, quality control, sequencing, and statistical analysis were carried out by Novogene Co., Ltd. Raw data of fastq format were firstly processed through fastp software. In this step, clean data (clean reads) were obtained by removing reads containing adapter, reads containing ploy-N, and low-quality reads from raw data. HISAT2 (version 2.2.1 (Mortazavi et al., 2008)) was used to align paired-end clean reads to the reference genome. The mapped reads of each sample were assembled by StringTie (version 3.0.1 (Pertea et al., 2015)) in a reference-based approach.

To estimate gene expression levels, featureCounts (version 2.2.1) was first used to count the reads mapped to each gene. The expected number of Fragments Per Kilobase of transcript sequence per Million base pairs sequenced (FPKM) was calculated based on the length of each gene and reads count mapped to each gene. Differential expression analysis was performed using the DESeq2 R package (version 1.42.0). The resulting p value is adjusted using the Benjamini and Hochberg methods to control FDR.

### Pathways analysis

Gene set enrichment analysis (https://doi.org/10.1073/pnas.0506580102) was used to calculate significantly over-/under-enriched Hallmark (Liberzon et al., 2015) and Reactome (database release #96 (Ragueneau et al., 2026)) gene sets satisfying an FDR-corrected p value < 0.05. See (Subramanian et al., 2005) for description of the methods used to generate normalised enrichment scores and p value statistics.

### Analysis of proteomics data

Raw mass-spectrometry data were analysed in FragPipe (version 23.1). Samples were searched against the UniProt Equus caballus reference proteome (UP000002281; downloaded in April 2026, containing 69,432 entries), including contaminants and decoys. For database searching and quantification, the predefined SILAC3 workflow was used with minor adjustments to default parameters. For MSFragger searches, MS1 and MS2 mass tolerances were set to 10 ppm and 20 ppm, respectively. Strict trypsin specificity with up to two missed cleavages was applied. Variable modifications included hydroxylation (P), heavy lysine (K6.025107), oxidation (M), and N-terminal acetylation; carbamidomethylation (C) was set as a fixed modification.

For MS1-based quantification, K0 was defined as the light channel and K6.020129 as the heavy channel. Search results were further processed in Perseus, version 2.0.7.0. Intensity values were log₂ transformed, normalised by median subtraction within samples, and filtered based on data completeness (≥3 valid values in at least one biological group). Missing values were imputed from a down shifted normal distribution. Differential abundance analysis was performed using a two sample Student’s t test combined with permutation-based FDR control (FDR set to 5 %).

Data was filtered for matrix proteins using the Matrisome database (Shao et al., 2023) and protein half-lives were calculated by comparing the abundances of heavy- and light-labelled proteins across time points, adapting previous methods to account for the reduced serum growth condition (Pickard et al., 2025; Schwanhäusser et al., 2009). For each biological subject and time point, the heavy:light ratio (*^H^*⁄*_L_*) of each detected matrisome protein was calculated. Rate constants of protein turnover, *k*, for each biological subject were calculated across 48 hour and 72 hour time points,

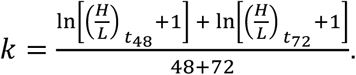

Where protein turnover rate constants could be calculated for at least 2 of 4 Lo-P and Hi-P biological subjects, turnover rate constants were then averaged across biological subjects and used to determine protein half-life,

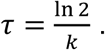

The mass spectrometry raw data included in this paper had been deposited to the Proteome eXchange Consortium via the PRIDE partner repository with the dataset identifier PXD080600 (Perez-Riverol et al., 2025).

## Statistical analyses

All statistical analyses were performed using Prism software (version 10.2.3, GraphPad). Unless indicated otherwise, data are represented as mean ± SD, and p values show the results of a paired two-tailed t test (∗, < 0.05; ∗∗, < 0.01; ∗∗∗, < 0.001, ∗∗∗∗, < 0.0001). The number of replicates represents the number of samples from different animals used.

## Resources table

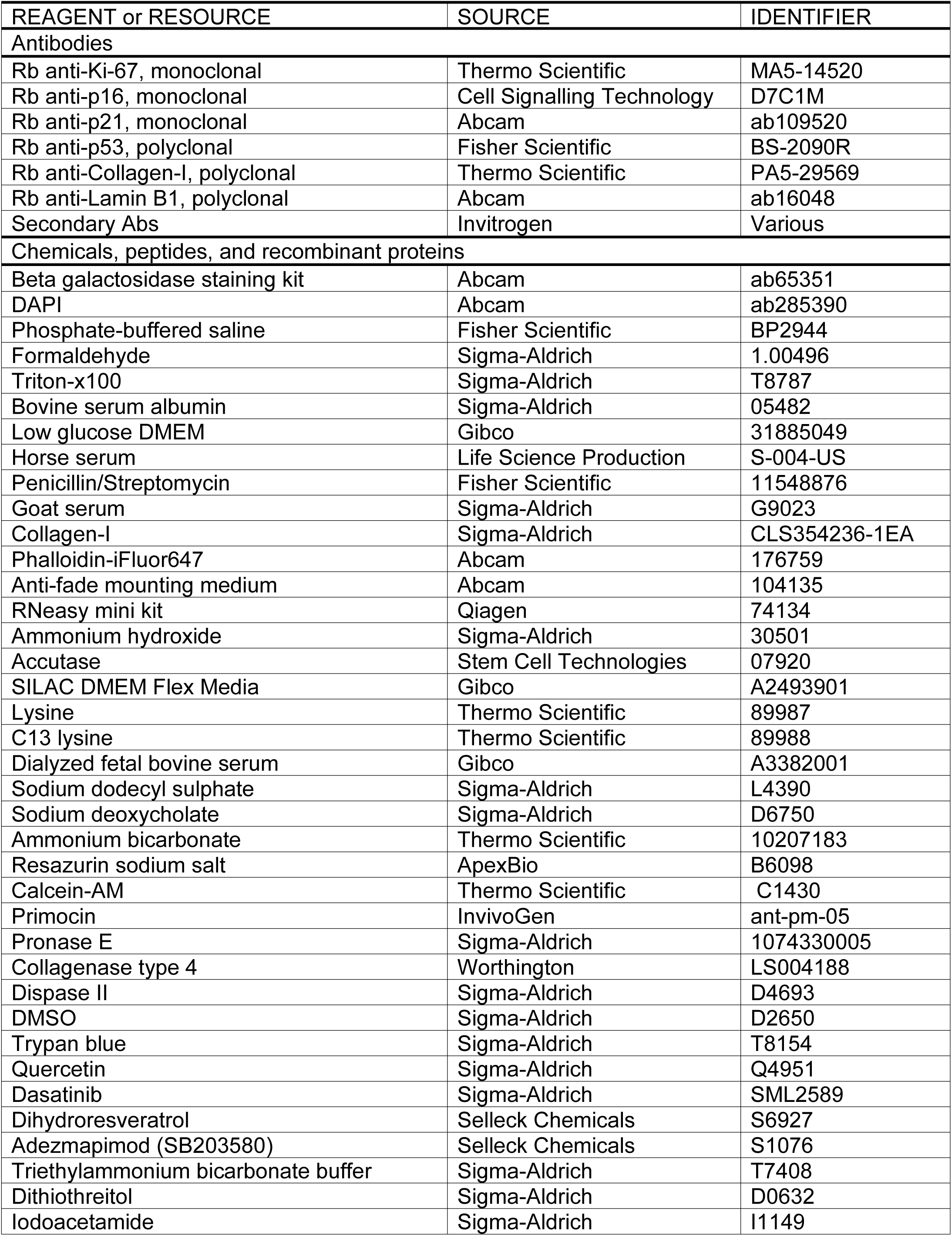
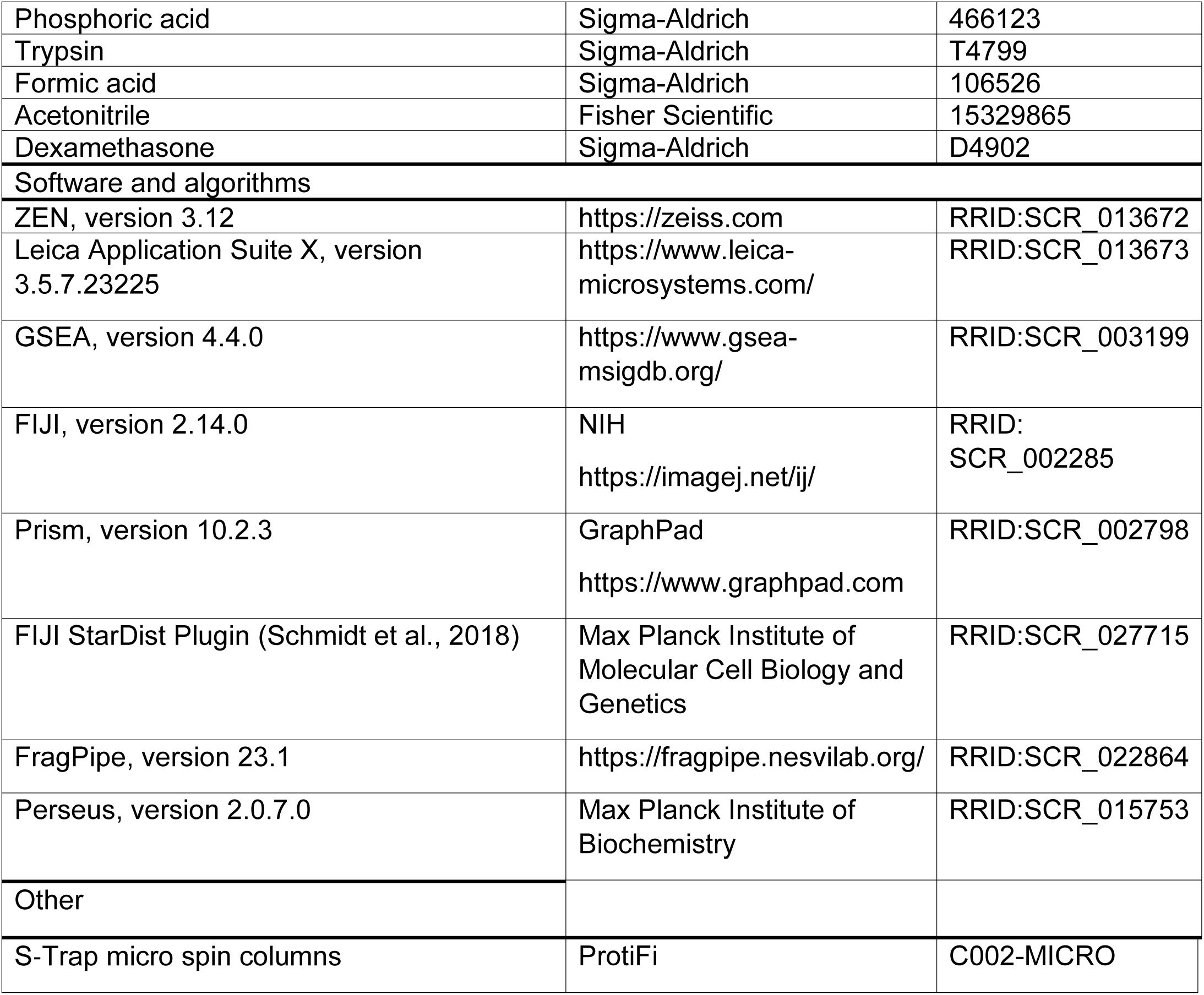

| REAGENT or RESOURCE | SOURCE | IDENTIFIER |
| --- | --- | --- |
| Antibodies |  |  |
| Rb anti-Ki-67, monoclonal | Thermo Scientific | MA5-14520 |
| Rb anti-p16, monoclonal | Cell Signalling Technology | D7C1M |
| Rb anti-p21, monoclonal | Abcam | ab109520 |
| Rb anti-p53, polyclonal | Fisher Scientific | BS-2090R |
| Rb anti-Collagen-I, polyclonal | Thermo Scientific | PA5-29569 |
| Rb anti-Lamin B1, polyclonal | Abcam | ab16048 |
| Secondary Abs | Invitrogen | Various |
| Chemicals, peptides, and recombinant proteins |  |  |
| Beta galactosidase staining kit | Abcam | ab65351 |
| DAPI | Abcam | ab285390 |
| Phosphate-buffered saline | Fisher Scientific | BP2944 |
| Formaldehyde | Sigma-Aldrich | 1.00496 |
| Triton-x100 | Sigma-Aldrich | T8787 |
| Bovine serum albumin | Sigma-Aldrich | 05482 |
| Low glucose DMEM | Gibco | 31885049 |
| Horse serum | Life Science Production | S-004-US |
| Penicillin/Streptomycin | Fisher Scientific | 11548876 |
| Goat serum | Sigma-Aldrich | G9023 |
| Collagen-I | Sigma-Aldrich | CLS354236-1EA |
| Phalloidin-iFluor647 | Abcam | 176759 |
| Anti-fade mounting medium | Abcam | 104135 |
| RNeasy mini kit | Qiagen | 74134 |
| Ammonium hydroxide | Sigma-Aldrich | 30501 |
| Accutase | Stem Cell Technologies | 07920 |
| SILAC DMEM Flex Media | Gibco | A2493901 |
| Lysine | Thermo Scientific | 89987 |
| C13 lysine | Thermo Scientific | 89988 |
| Dialyzed fetal bovine serum | Gibco | A3382001 |
| Sodium dodecyl sulphate | Sigma-Aldrich | L4390 |
| Sodium deoxycholate | Sigma-Aldrich | D6750 |
| Ammonium bicarbonate | Thermo Scientific | 10207183 |
| Resazurin sodium salt | ApexBio | B6098 |
| Calcein-AM | Thermo Scientific | C1430 |
| Primocin | InvivoGen | ant-pm-05 |
| Pronase E | Sigma-Aldrich | 1074330005 |
| Collagenase type 4 | Worthington | LS004188 |
| Dispase II | Sigma-Aldrich | D4693 |
| DMSO | Sigma-Aldrich | D2650 |
| Trypan blue | Sigma-Aldrich | T8154 |
| Quercetin | Sigma-Aldrich | Q4951 |
| Dasatinib | Sigma-Aldrich | SML2589 |
| Dihydroresveratrol | Selleck Chemicals | S6927 |
| Adezmapimod (SB203580) | Selleck Chemicals | S1076 |
| Triethylammonium bicarbonate buffer | Sigma-Aldrich | T7408 |
| Dithiothreitol | Sigma-Aldrich | D0632 |
| Iodoacetamide | Sigma-Aldrich | I1149 |
| Phosphoric acid | Sigma-Aldrich | 466123 |
| Trypsin | Sigma-Aldrich | T4799 |
| Formic acid | Sigma-Aldrich | 106526 |
| Acetonitrile | Fisher Scientific | 15329865 |
| Dexamethasone | Sigma-Aldrich | D4902 |
| Software and algorithms |  |  |
| ZEN, version 3.12 | https://zeiss.com | RRID:SCR_013672 |
| Leica Application Suite X, version 3.5.7.23225 | https://www.leica-microsystems.com/ | RRID:SCR_013673 |
| GSEA, version 4.4.0 | https://www.gsea-msigdb.org/ | RRID:SCR_003199 |
| FIJI, version 2.14.0 | NIH<br>https://imagej.net/ij/ | RRID:SCR_002285 |
| Prism, version 10.2.3 | GraphPad<br>https://www.graphpad.com | RRID:SCR_002798 |
| FIJI StarDist Plugin (Schmidt et al., 2018) | Max Planck Institute of Molecular Cell Biology and Genetics | RRID:SCR_027715 |
| FragPipe, version 23.1 | https://fragpipe.nesvilab.org/ | RRID:SCR_022864 |
| Perseus, version 2.0.7.0 | Max Planck Institute of Biochemistry | RRID:SCR_015753 |
| Other |  |  |
| S-Trap micro spin columns | ProtiFi | C002-MICRO |

## Acknowledgements

JL, RKS, JD, RGAF, and CT were funded to perform the study by the Biotechnology and Biological Sciences Research Council (BBSRC, BB/W007282/1). NI was supported by funding from the Horserace Betting Levy Board (HBLB, vet/prj/789). Biological samples were provided by F Drury and Sons Ltd, and Nicole Larkin, who was supported by funding from the Hong Kong Jockey Club (HKJC, MRG-241027 URN 2024 2343- A). RNA sequencing was carried out by Novogene Co., Ltd.

## Author contributions

JL, RKS, JD, RF, and CT designed research. JL, NI, KC, and HD performed research. AH, IV, and GB provided mass spectrometry expertise and technical services. JL and CT wrote the paper.

## Supplemental Information

### Supplemental Figures

**Figure S1.**
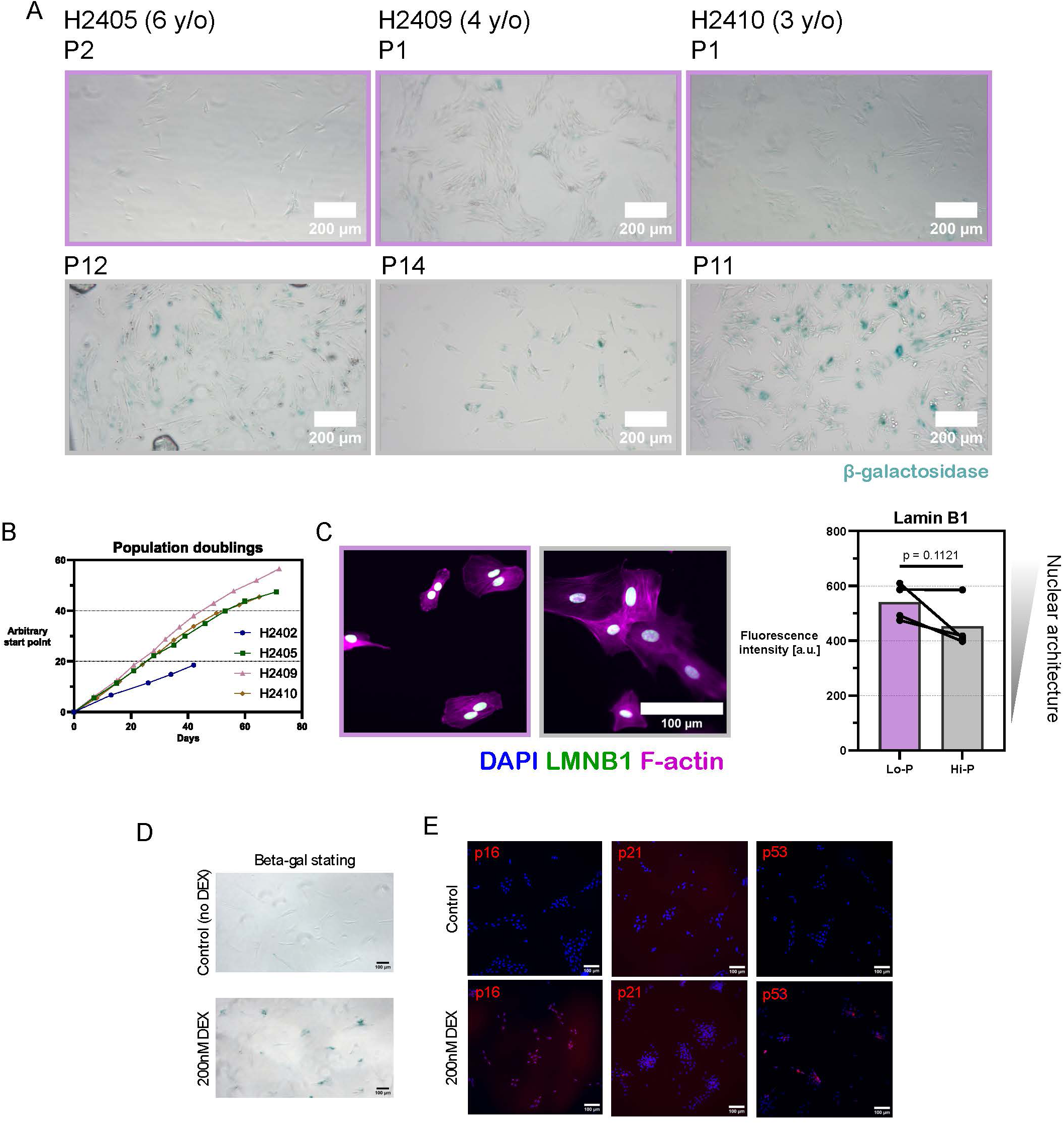
Validation of cellular senescence in high-passage tenocyte cultures. **(A)** Supplemental images from three additional biological replicates of low- (Lo-P, purple) and high-passage (Hi-P, grey) cells stained for senescence-associated beta-galactosidase. **(B)** Population doublings of individual biological replicates over time, with points indicating passages. Recording of population doublings began (0,0) at an arbitrary time and passage # for each replicate. Repeated passaging was ended at the point that further passaging would not yield enough cells for downstream experiments. **(C)** Immunofluorescence of Lamin B1 protein in Lo-P and Hi-P tenocytes. Arbitrary units. N = 4. **(D,E)** The phenotype induced by replicative stress mirrors the senescence phenotype induced by dexamethasone treatment, with upregulation of (C) β-galactosidase; (D) p16, p21, and p53. Unless otherwise specified, data are represented as mean ± SD, and p values show the results of a paired two-tailed t test.

**Figure S2.**
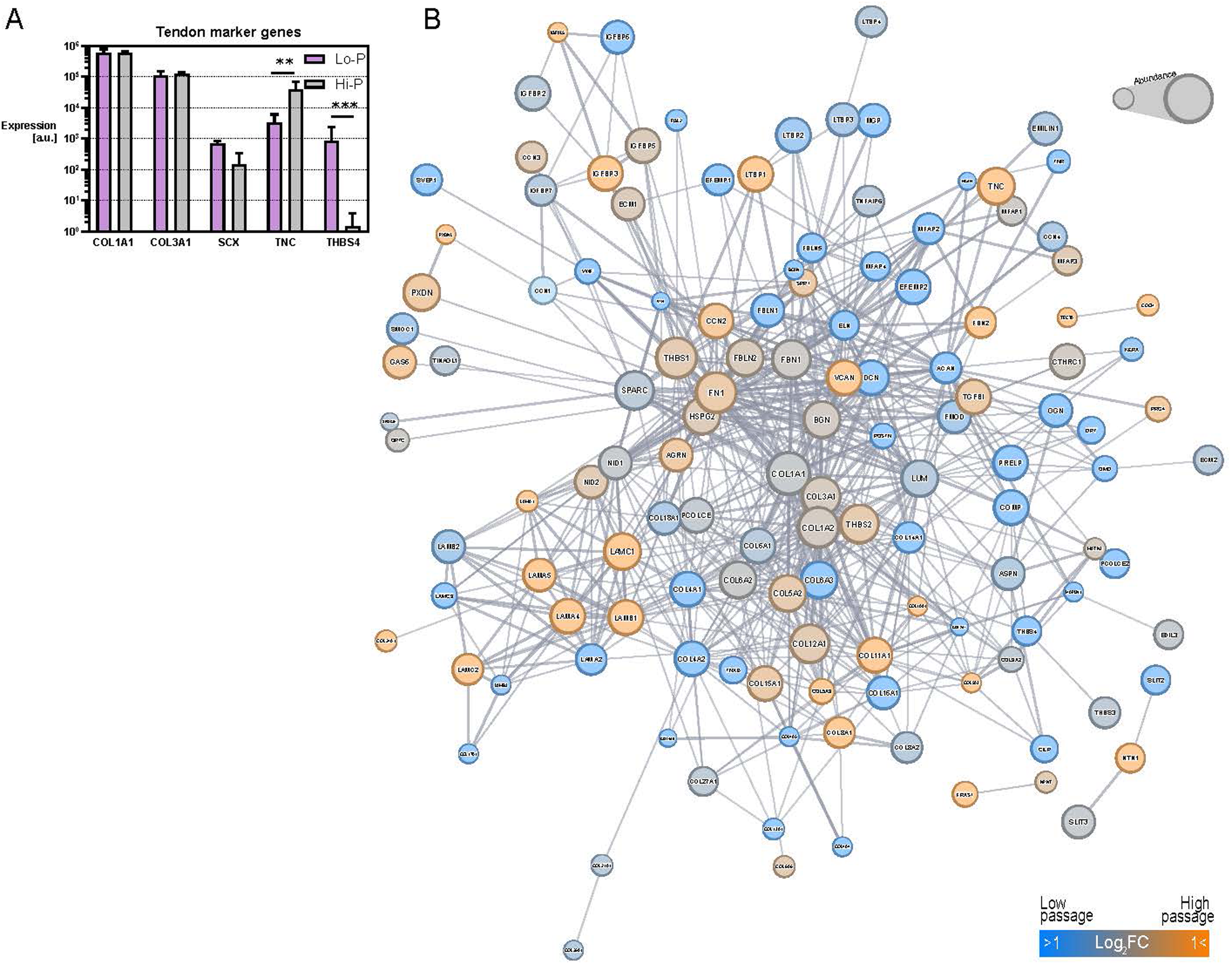
Extended transcriptomics of high-passage tenocytes. **(A)** Transcriptional changes in tendon-specific markers between low- (Lo-P) and high-passage (Hi-P) cells. N = 4. Data are represented as mean ± SD, and p values from RNA sequencing are calculated as described in Methods (∗∗ < 0.01, ∗∗∗ < 0.001). **(B)** Interaction map with nodes consisting of transcripts corresponding to matrisome proteins. Nodes are sized according to their signal strength in Lo-P cells in our sequencing data, coloured according to their fold-change in high passage cells, and edges are weighted by StringDB interaction score, showing only edges which satisfy the highest confidence filter.

**Figure S3.**
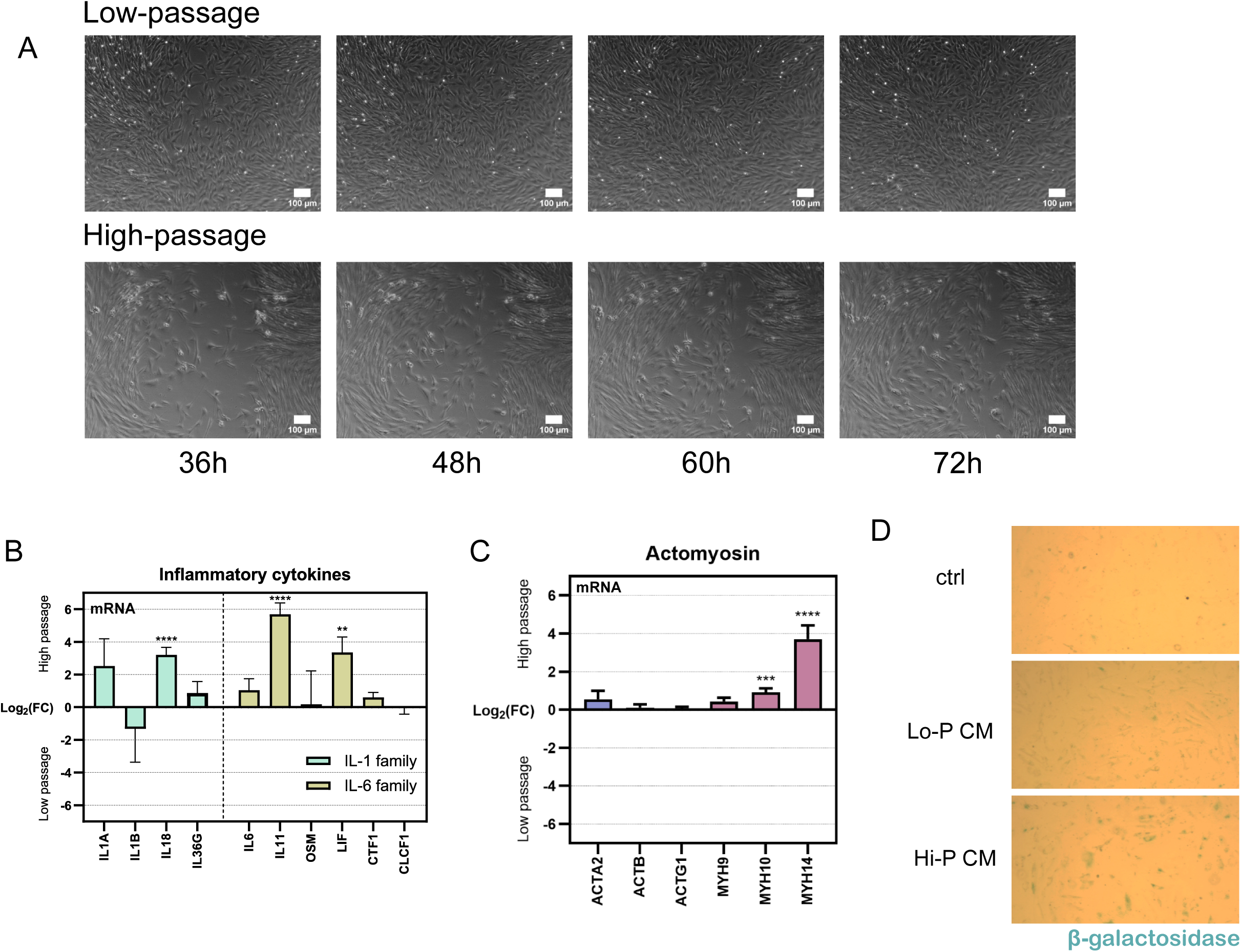
Extended functional consequences of tenocyte senescence. **(A)** Scratch assay showing live microscopy images of low- (Lo-P) and high-passage (Hi-P) cultures 36-72 hours following introduction of an artificial would site. N = 4. **(B)** Transcriptional changes in IL-1 and IL-6 family cytokines between Lo-P and Hi-P tenocytes. N = 4. **(C)** Transcriptional changes in actomyosin machinery between Lo-P and Hi-P tenocytes. N = 4. **(D)** Lo-P tenocytes treated for four days with either Lo-P or Hi-P conditioned media or control media and stained for the senescence marker β-galactosidase. Data are represented as mean ± SD, and p values from RNA sequencing are calculated as described in Methods (∗∗ < 0.01, ∗∗∗ < 0.001, ∗∗∗∗ < 0.0001).

**Figure S4.**
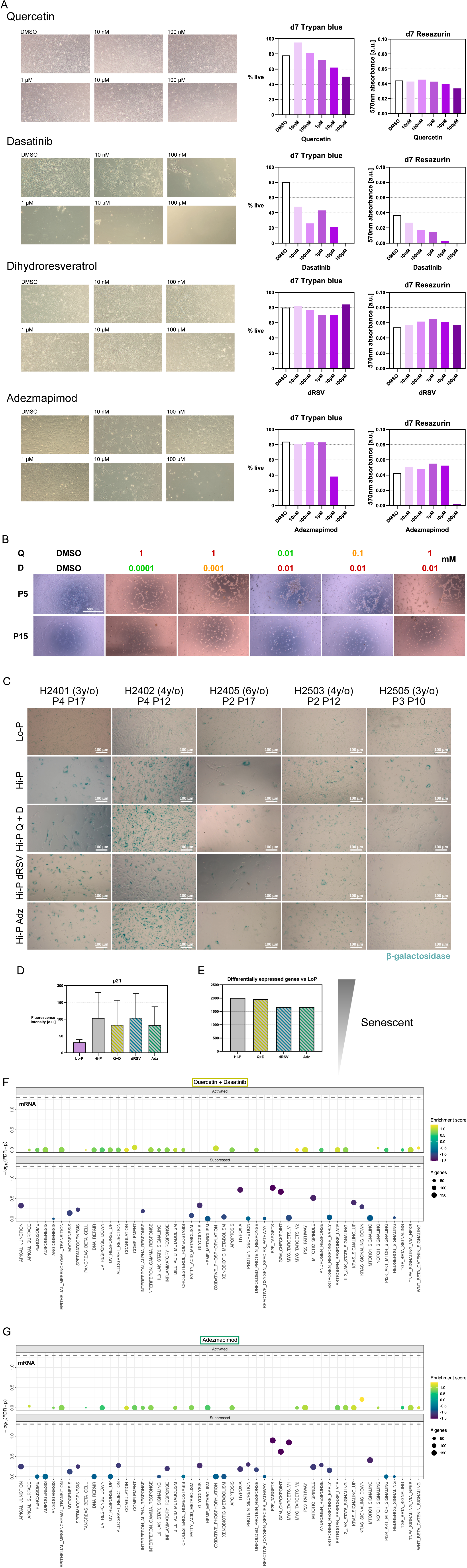
Characterisation of the effects of senotherapeutics on equine tenocytes. **(A)** Toxicity assays of senotherapeutic compounds comparing cell growth (bright field images), cell viability (trypan blue) and cell metabolism (570 nm resazurin absorbance) in low-passage cells after one week of culture with various dosages. N = 1. **(B)** Bright field images of trials of 24-hour dosages of quercetin (Q) and dasatinib (D) on low- (Lo-P) and high-passage (Hi-P) cells. N = 1. **(C)** Supplemental bright field images of cells stained for senescence-associated β-galactosidase after seven days treatment + three days recovery in Lo-P and Hi-P tenocytes, plus Hi-P tenocytes treated with senolytics (Q+D), senoreverter (dRSV), or senomorphic (Adz). N = 5. **(D)** Quantification of p21 immunofluorescence after seven days treatment + three days recovery in Hi-P tenocytes treated with senolytics (Q+D), senoreverter (dRSV), or senomorphic (Adz). Data are represented as mean ± SD, N = 6. **(E)** Number of differentially expressed genes between the transcriptome of Lo-P tenocytes and the transcriptomes of Hi-P tenocytes either vehicle-only or treated with senolytics (Q+D), senoreverter (dRSV), or senomorphic (Adz). **(F,G)** Gene Set Enrichment Analysis (GSEA) of enrichment in the Hallmark gene sets in high-passage tenocytes treated with (F) Q+D, and (G) Adz, versus vehicle-only Hi-P tenocytes. Dashed line indicates a false discovery rate-corrected p value < 0.05 (Subramanian et al., 2005). N = 3.

### Supplemental Videos

**Video S1**

72-hour wound-closure scratch assay of low-passage (Lo-P) cells, in reduced serum (0.5 %) media.

**Video S2**

72-hour wound-closure scratch assay of high-passage (Hi-P) cells, in reduced serum (0.5 %) media.

